# Using embedded alginate microparticles to tune the properties of *in situ* forming poly(*N*-isopropylacrylamide)-graft-chondroitin sulfate bioadhesive hydrogels for replacement and repair of the nucleus pulposus of the intervertebral disc

**DOI:** 10.1101/2021.04.16.439319

**Authors:** Thomas Christiani, Karen Mys, Karl Dyer, Jennifer Kadlowec, Cristina Iftode, Andrea Jennifer Vernengo

**Affiliations:** Rowan University, Department of Biomedical Engineering, 201 Mullica Hill Rd, Glassboro, NJ 08028, USA; AO Research Institute Davos Clavadelerstrasse 8, Davos Platz 7270 Switzerland; Rowan University, Department of Mechanical Engineering, 201 Mullica Hill Rd, Glassboro, NJ 08028, USA; Baldwin Wallace University, Department of Computer Science and Engineering 275 Eastland Rd, Berea, OH 44017, USA; Rowan University, Department of Molecular and Cellular Biosciences, 201 Mullica Hill Rd, Glassboro, NJ 08028, USA; Rowan University, Department of Chemical Engineering, 201 Mullica Hill Rd, Glassboro, NJ 08028, USA

## Abstract

Low back pain (LBP) is a major public health issue associated with degeneration of the intervertebral disc (IVD). The early stages of degeneration are characterized by the dehydration of the central, gelatinous portion of the IVD, the nucleus pulposus (NP). One possible treatment approach is to replace the NP in the early stages of IVD degeneration with a hydrogel that restores healthy biomechanics while supporting tissue regeneration. The present study evaluates a novel thermosensitive hydrogel based on poly(*N*-isopropylacrylamide-graft-chondroitin sulfate) (PNIPAAM-g-CS) for NP replacement. The hypothesis was tested that the addition of freeze-dried, calcium crosslinked alginate microparticles (MPs) to aqueous solutions of PNIPAAm-g-CS would enable tuning of the rheological properties of the injectable solution, as well as the bioadhesive and mechanical properties of the thermally precipitated composite gel. Further, we hypothesized that the composite would support encapsulated cell viability and differentiation. Structure-material property relationships were evaluated by varying MP concentration and diameter. The addition of high concentrations (50 mg/mL) of small MPs (20 ± 6 µm) resulted in the greatest improvement in injectability, compressive mechanical properties, and bioadhesive strength of PNIPAAm-g-CS. This combination of PNIPAAM-g-CS and alginate MPs supported the survival, proliferation, and differentiation of adipose derived mesenchymal stem cells (ADMSCs) towards an NP-like phenotype in the presence of soluble GDF-6. When implanted *ex vivo* into the intradiscal cavity of degenerated porcine IVDs, the formulation restored the compressive and neutral zone (NZ) stiffnesses to intact values and resisted expulsion under lateral bending. Overall, results indicate the potential of the hydrogel composite to serve as a scaffold for supporting NP regeneration. This work uniquely demonstrates that encapsulation of re-hydrating polysaccharide-based MPs may be an effective method for improving key functional properties of *in situ* forming hydrogels for orthopaedic tissue engineering applications.

## 1. Introduction

Low back pain (LBP) is a ubiquitous public health issue which burdens health care systems world-wide, affecting up to 80% of adults [1, 2]. LBP in patients is often associated with degeneration of the intervertebral disc (IVD) [3-6]. The intervertebral disc (IVD) is the load bearing joint between vertebrae consisting of a central nucleus pulposus (NP) and peripheral annulus fibrosus (AF). The NP is an amorphous gel comprised of collagen II and elastin fibers dispersed in a water-rich aggrecan phase [7]. In contrast, the AF has an organized, anisotropic structure made up of lamellae, or multilayered, oriented collagen fibers in an angle-ply structure [8]. The highly swellable NP expands radially under compressive loads and transfers the loads to the outer AF in circumferential tension [9]. With aging, increased catabolism reduces the collagen II and aggrecan content of the NP [10], resulting in loss of its swelling capacity, a change in the load distribution, and the formation of tears and fissures in the AF. These structural changes may be accompanied by vascularization and neoinnervation [11], associated with pain [12, 13]. Current clinical treatments for low back pain include a combination of analgesics with physical therapy or discectomy to remove nerve impinging disc tissue [14]. While these interventions provide immediate pain relief, they do not restore the healthy biomechanics to the tissue, so degeneration can continue or even accelerate. [15-17]

Because early stage IVD degeneration is characterized primarily by changes in the NP region, the tissue is a target for newly developed therapeutic interventions. For instance, if the annulus and endplates are still competent, a swellable biomaterial can be implanted to replace the dehydrating NP. Injectability is considered paramount so the hydrogel can be implanted intradiscally with minimal damage to the AF and fill irregularly shaped tissue defects in the NP. Due to high *in vivo* intradiscal pressures [18], the injectable hydrogel solution must have sufficient viscosity to be injected into an NP region without extravasation [19]. Once cured, the hydrogel should deform mechanically like the native healthy NP [20, 21]. For tissue engineering approaches to repairing the IVD, the injectable hydrogel must meet these requirements and also support viability of encapsulated cells and prevent their leakage during motion and loading [22, 23]. Towards these design goals, multiple *in situ* forming hydrogel materials have been studied for NP replacement and regeneration, like decellularized matrix-based systems [24, 25], alginate [26], collagen [27, 28], hyaluronic acid [29, 30], and chitosan [31].

Notably, in an *ex vivo* test with ovine IVDs [20], neither the implantation of hydrogel NP replacements or the re-implantation of the natural nucleus tissue restored functionality of an intact disc. It was concluded that integration with the surrounding AF tissue is a critical component of an NP replacement strategy. Thus, in parallel with the development of injectable cell-friendly systems with tunable mechanical properties, recent research has focused on engineering hydrogel bioadhesives that adhere to surrounding tissue in the IVD to minimize the risk of herniation and improve biomechanical performance. Fibrin is a biocompatible, *in situ*-forming carrier that forms an adhesive bond with tissue [32]. However, fibrin degrades quickly, thus making it non-ideal for the long repair process of the IVD [33]. The low mechanical properties [34, 35] and bioadhesive strength [36-38] make it inappropriate for load bearing applications. Genipin-crosslinked fibrin has been investigated as a bioadhesive cell carrier for AF repair. The covalent crosslinking of fibrin with genipin improves mechanical stiffness and adhesive strength of fibrin [39], but genipin can have potentially cytotoxic effects [40-42]. An inherent challenge with fibrin-genipin is to balance the composition to improve the material properties of fibrin while promoting cell survival and extracellular matrix (ECM) deposition [43]. Injectable hydrogels based on tyramine-modified hyaluronic acid hydrogels crosslinked with horseradish peroxidase (HRP) and hydrogen peroxide (H_2_O_2_) were investigated as a bioadhesive cell carrier. However, the bonding strength to cartilage was not statistically significantly different than fibrin glue [44].

In response to these needs in NP repair, we sought to design a novel *in situ* forming cell carrier for NP replacement with bioadhesive properties. Previously, we reported on a novel thermosensitive graft copolymer, poly(*N*-isopropylacrylamide)-graft-chondroitin sulfate (PNIPAAm-g-CS) [45, 46]. Aqueous solutions of PNIPAAm-g-CS behave as a hydrophilic, flowable liquid below the lower critical solution temperature (LCST) of approximately 30°C, and a precipitated, soft hydrogel above the LCST. Due to this phase transition, the copolymer can be injected into the intradiscal cavity through a small gauge needle and form a space-filling gel *in situ* which is compatible with encapsulated cells [45]. The limitations of using PNIPAAm-g-CS for NP replacement are its low solution viscosity below the LCST and limited bioadhesive properties, which allow immediate extravasation upon injection into an intradiscal cavity. Therefore, we sought to improve these properties of PNIPAAm-g-CS for NP replacement by generating a composite with calcium crosslinked alginate microparticles (MPs).

Multiple levels of rationale were used for tuning PNIPAAm-g-CS properties with MPs. There is an established link between particle-scale motion and rheological and mechanical properties of a suspension [47-49]. The viscosity of a solution increases with the addition of fine particles due to increased packing density and molecular interactions during deformation [50]. Viscosity is an important parameter for mediating adherence to tissue, since polymeric solutions that are too liquid-like lack the cohesion necessary to form substantial interactions with the tissue [51]. Embedding MPs in a hydrogel increases surface roughness [52], which can promote mechanical fixation with a tissue substrate. Further, MPs incorporated into bulk hydrogel structures enhance network toughness [53, 54]. DeVolder et. al. showed that poly(lactic-co-glycolic acid) MP incorporation into 3D crosslinked collagen networks increased stiffness and elasticity [55]. Notably, the encapsulation of MPs within hydrogel networks has been reported to improve mechanical performance while preserving encapsulated cell viability [56, 57]. In the present study, calcium crosslinked alginate was selected to comprise the MPs because the material is inexpensive, biocompatible, hydrophilic, and can be fabricated without the use of toxic reagents. Alginate has an abundance of hydrophilic hydroxyl and carboxylic acid groups, as well as a net anionic charge [58], facilitating swellability and physical interaction with proteins in the extracellular matrix (ECM) [59, 60].

Herein, we test the hypothesis that the addition of calcium crosslinked alginate MPs to aqueous solutions of PNIPAAm-g-CS would enable tuning of the rheological properties of the injectable solution, as well as the bioadhesive and mechanical properties of the precipitated gel composite, improving the suitability of the material for NP replacement. Further, we hypothesized that the composite would support encapsulated cell viability, NP differentiation, and ECM synthesis.

This study was comprised of four aims: 1) To study structure-property relationships in injectable PNPAAm-g-CS + MP composites by varying MP concentration and diameter. Subsequently, we aimed to evaluate the formulation most closely mimicking the native NP for its ability to 2) support NP regeneration *in vitro* by encapsulated adipose derived mesenchymal stem cells (ADMSCs), 3) restore the degenerated porcine IVD compressive biomechanical properties *ex vivo*, and 4) resist expulsion from the porcine IVD cavity under lateral bending.

## 2. Materials and Methods

### 2.1 Graft Copolymer Synthesis

Poly(*N*-isopropylacrylamide)-graft-chondroitin sulfate (PNIPAAm-g-CS) was synthesized using free radical polymerization of N-isopropylacrylamide (NIPAAm) and methacrylated chondroitin sulfate (mCS) as described in previous publications [45, 46]. Based on previous results, a copolymer with a molar ratio of 1000:1 (NIPAAm:mCS) and mCS with a methacrylate degree of substitution of 0.1 was used [45, 46].

### 2.2 Microparticle Synthesis

A water-in-oil emulsion technique was used to synthesize alginate MPs of varying diameters as described in previous publications [45, 46]. Briefly, 2 % (w/v) alginate solution (Sigma-Aldrich) and 1 % (v/v) Tween 20 surfactant (Sigma-Aldrich) were emulsified in a vegetable oil phase. Low and high stir speeds were used to alter alginate and oil droplet size, resulting in large and small MP diameters, respectively. A 2 % (w/v) CaCl_2_ solution was added dropwise to the emulsion to crosslink the alginate. Residual oil was removed from crosslinked MPs through a series of alternating centrifugation (500 x g for 5 minutes) and washing steps using 70 % (v/v) isopropanol and deionized water. An average size was calculated for each batch by measuring the diameters of 50 randomly selected MPs. Alginate MPs were freeze dried and stored at 4 °C until further use.

### 2.3 Composite Preparation and Factorial Design

Freeze dried PNIPAAm-g-CS was dissolved in 0.01 M PBS at a concentration of 5 % (w/v) and blended with freeze-dried alginate MPs to create the composite hydrogels. The same batches of hydrogel and alginate MPs were used for each individual study. Batch consistency between studies was maintained by monitoring hydrogel viscosity and MP diameter. A 2×2 factorial design was used to study the effects of small (S, 20.0 ± 6.0 μm) and large (L, 120.0 ± 39 μm) MPs and low (25 mg/mL) and high (50 mg/mL) MP concentrations on scaffold properties. Results for four different PNIPAAm-g-CS + MP formulations, S-25, L-25, S-50, L-50 were compared to PNIPAAM-g-CS alone (P-0). Sample compositions are summarized in **Table 1**. The compositions were selected based on preliminary studies [61] demonstrating that lower MP concentrations (below 25 mg/mL) and higher MP diameters (above 150 μm) produced less favourable impacts on PNIPAAm-g-CS bioadhesive strength.

**Table 1.** Formulations of 5 % (w/v) PNIPAAm-g-CS with or without suspended alginate MPs of various concentrations and diameters were evaluated in this study.

| Formulation Designation | MP Diameter (μm) | MP Concentration (mg/mL) |
| --- | --- | --- |
| P-0 | n/a | 0 |
| S-25 | 20.0 ± 6.0 | 25 |
| S-50 | 20.0 ± 6.0 | 50 |
| L-25 | 120.0 ± 39 | 25 |
| L-50 | 120.0 ± 39 | 50 |

### 2.4 Characterization of material properties

#### 2.4.1 Swelling Properties

Approximately 500 µL of each solution formulation (*n* = 5 per group) was gelled at 37 °C and swelled *in vitro* in 0.01 M PBS for 14 days. The PBS solutions were refreshed every other day. The swelling ratio for each sample at the beginning and end of the study was calculated as the wet weight divided by the dry weight.

#### 2.4.2 Scanning Electron Microscopy

Scaffold architecture was evaluated over the 14-day swelling period using a Phenom Pure scanning electron microscope (SEM) (Nanoscience Instruments) equipped with a cryostage. Immediately prior to analysis, the gel samples were removed from PBS, directly placed on pre-warmed foil wraps, flash frozen in liquid nitrogen, and imaged at – 20 °C.

#### 2.4.3 Enzymatic Degradation

Approximately 0.3 mL of samples P-0 or S-50 (*n* = 5 per group) was immersed in 2 mL of 0.01 M PBS containing either 0.1 mg/mL collagenase P, 0.01 mg/mL hyaluronidase, 50 ng/mL aggrecanase, or 0.1 U/mL chondroitinase ABC (Sigma Aldrich). Enzyme solution was maintained at 37 °C and refreshed each day for 7 days. As a control, formulations were exposed to 0.01 M PBS without enzyme. The percent mass loss was calculated using Equation 1:

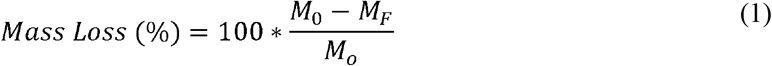

where, M_0_ and M_F_ are the initial and final dry masses of the sample, respectively.

#### 2.4.4 Rheological Characterization

The rheological properties of each formulation were characterized using a Texas Instrument DHR-3 rheometer. A 20 mm parallel plate configuration with a 500 µm gap (160 µL sample volume) was used for each test (*n* = 5). Temperature ramps were performed within the range of 25 to 37 °C at 1 °C/min and a constant 1 % strain and 1 Hz frequency. Gel points were identified as the crossover of the storage modulus (G’) and loss modulus (G’’). Frequency sweeps were performed within the range of 0.01 to 15 Hz with a constant 1 % strain and temperature of 37 °C. The parameters G’, G’’, complex modulus (G*), and phase shift angle (δ) were quantified for each formulation.

#### 2.4.5 Bioadhesive Properties

All *in vitro* adhesive and mechanical characterization studies were performed on a Shimpo E-Force Test Stand with a 2 N load cell (FGV-0.5XY). Tensile and lap shear tests were executed based on ASTM standards F2258-05 and F2255-05, respectively. For the tensile tests, 25 µL of cold hydrogel solution was pipetted between porcine inner AF tissue substrates, which were cut to 0.5 cm^2^ and spaced 1mm apart. Hydrogel-substrate combinations were immersed in a 37 °C water bath for 5 minutes under a preload of 0.01 N before application of the tensile strain at a rate of 5 mm/min.

For the lap shear tests, 50 µL of hydrogel solution was applied between rectangular substrates (0.5 cm x 1 cm), equilibrated for 5 minutes of gelation at 37 °C in the water bath, then sheared at a rate of 5 mm/min. The ultimate adhesive tensile and shear strengths were determined from the load-displacement data normalized to the cross-sectional area of the AF tissue. To visualize biomaterial interaction with the substrates, a set of samples were collected for histological assessment immediately after application to the tissue. The tissue-biomaterial constructs were fixed with 4% formaldehyde in PBS for 24 h at 37 °C, embedded in frozen section compound, sectioned to 30 µm, and stained with alcian blue to enable visualization of the tissue, rich in glycosaminoglycan (GAG), and the biomaterials, which non-specifically absorb the dye. The cells in the tissue were counterstained with Weigert’s hematoxylin.

#### 2.4.6 Unconfined Compressive Properties

For the unconfined compression tests (*n* = 5), cylindrical hydrogels (formulations P-0, S-25, S-50, L-25, and L-50) were pre-formed in 48-well plates at 37 °C for 5 minutes. Then, the hydrogels were transferred to a 37 °C saline bath where they were deformed at a rate of 1 mm/min. Data were normalized to stress and strain using the initial cross-sectional area and height of each hydrogel and the unconfined compressive moduli reported at 25% strain.

#### 2.4.7 Confined Compressive Properties

Confined compression tests (*n* = 7) were performed based on ASTM F2789-10 using a custom-built apparatus with a surrogate AF mold composed of RTV-630 silicone elastomer (Momentive Performance Materials Inc.). The apparatus was encased in plexiglass filled with saline maintained at 37 °C. Approximately 350 µL of hydrogel solution was injected into the mold and allowed to equilibrate to temperature before deforming at a rate of 1 mm/min. Data were normalized to stress and strain using the initial cross-sectional area and height of each hydrogel and the confined compressive moduli reported at 25% strain.

Due to its bioadhesive properties, ease of injectability, and mechanical performance approaching the native NP, formulation S-50 was the focus of subsequent *in vitro* cell culture studies and *ex vivo* biomechanical tests. As a control for the cell viability analysis, metabolic activity assay, and histological characterization, cell encapsulation within P-0 was evaluated in parallel.

### 2.5 *In vitro* cell culture studies

#### 2.5.1 Expansion of ADMSCs

Commercial normal human ADMSCs (ScienCell, female donor, 30 years old) were expanded in monolayers using MSC basal medium (ScienCell) containing 5 % fetal bovine serum (FBS), 5 % MSC growth supplement, and 5 % penicillin/streptomycin solution and cultured in an incubator at 37 °C with 5 % CO_2_. ADMSCs were passaged to 80% confluency and used for all studies at passage 4.

#### 2.5.2 Hydrogel encapsulation of ADMSCs

Lyophilized PNIPAAm-g-CS and alginate MPs were sterilized by soaking in 70% isopropanol and exposure to UV light. Then, 5 % w/v PNIPAAm-g-CS was prepared in NP differentiation medium containing high glucose DMEM (Gibco), 1 % FBS (Gibco), insulin-transferrin-selenium-ethanolamine (ITS-X) (Gibco), 100 µM L-ascorbic acid-2-phosphate (Sigma-Aldrich), 1.25 mg/mL bovine serum albumin (BSA) (Sigma-Aldrich), 0.1 µM dexamethasone (Sigma-Aldrich), 40 µg/mL L-proline (Sigma-Aldrich), 5.4 µg/mL linoleic acid (Sigma-Aldrich), antibiotic-antimycotic (Gibco) and 100 ng/mL of GDF6 (PeproTech) [62]. Pelleted ADMSCs were combined with the solutions prepared in media of 5% PNIPAAM-g-CS + alginate MPs (S-50) or 5% PNIPAAm-g-CS (P-0). The final cell density in each of the formulations was 5 × 10^6^ cell/mL. Lastly, approximately 100 µL of each cell-seeded formulation was dispensed into a 48 well plate and gelled before adding 500 µL of NP differentiation media on top. Media was refreshed every other day and the formulations were cultured for 14 days.

#### 2.5.3 Evaluation of Cellular Viability and Metabolic Activity

Live/Dead™ Viability/Cytotoxicity Kit (Invitrogen) was used to assess ADMSC viability. At day 14 of culture, formulation S-50 or P-0 was dissolved in 0.01 M PBS containing 50 mM citrate (Sigma-Aldrich) and 20 mM EDTA (Sigma-Aldrich). Sodium citrate-EDTA buffer dilutes the hydrogel and reverses ionic alginate-Ca^2+^ crosslinks for complete removal of polymeric material, which obstructs visualization of the cells. Suspended cells were pelleted at 300 x g for 5 minutes at 4 °C and resuspended in Live/Dead™ reagent containing 2 µM calcein AM and 4 µM ethidium homodimer-1 in high glucose DMEM for 1 hour at 37 °C and 5 % CO_2_. Cells were isolated from the Live/Dead™ reagent, rinsed with 0.01 M PBS, dispensed in a 48 well plate, and imaged using an inverted fluorescent light microscope. Cellular viability was quantified using ImageJ software.

ADMSC metabolic activity was tracked over 14 days using the alamarBlue® Cell Viability Assay (Bio-Rad). Media was removed from samples of S-50 or P-0 (*n* = 5 each), replaced with 300 µL of 10 % alamarBlue reagent in NP differentiation medium, and incubated for 5 hours at 37 °C and 5 % CO_2_. Wells without cells were used to correct for background interference. Reduced reagents were removed from the samples and absorbance readings were measured using a spectrophotometer at 570 and 600 nm. Percent reagent reduction was calculated as described by the manufacturer’s instructions.

#### 2.5.4 Histology

Glycosaminoglycan (GAG) and collagen production were visualized histologically after 14 days of culture. Formulation P-0 or S-50 was fixed for 10 minutes with 4 % formaldehyde (Fisher Scientific), embedded in frozen section compound (VWR), snap-frozen in methylbutane chilled with liquid nitrogen, and sectioned to 20 µm sections. Since the polymers tend to absorb the histological dyes non-specifically, slides were rinsed with sodium citrate-EDTA buffer at room temperature to remove the PNIPAAM-g-CS and alginate MPs by dissolution. Then, GAG and collagen were stained using 1 % w/v alcian blue or 0.1 % w/v picrosirius red, respectively. Cell nuclei were counterstained with Weigert’s hematoxylin. ECM deposition was compared on days 0 and 14.

#### 2.5.5 Quantitative Polymerase Chain Reaction

Gene expression profiles of ADMSCs after 14 days of culture in formulation S-50 were examined using quantitative real-time polymerase chain reaction (qRT-PCR). Seeded cells were pelleted from PNIPAAm-g-CS + MPs gels by dissolution in sodium citrate-EDTA buffer and subsequent centrifugation. Total RNA was extracted using the Pure Link™ RNA Extraction Mini Kit (Ambion®, Life Technologies™), quantified in terms of concentration and purity with a nanodrop (Applied Biosystems), and reverse transcribed to cDNA using the High Capacity cDNA Reverse Transcription Kit (Applied Biosystems). Target genes (**Supplementary Table 1**) were amplified in 20 µL reactions using 20 ng of cDNA, Fast SYBR® Green Master Mix (Fisher Scientific), 500 nM primer concentrations, and an Applied Biosystems 9800 Fast Thermal Cycler. Relative gene expression was calculated using the delta-delta Ct method (2^-ΔΔCt^) and normalized to ADMSCs on day 0 after encapsulation and the housekeeping gene GAPDH.

#### 2.5.6 Immunofluorescence

For cells encapsulated in formulation S-50, an indirect immunofluorescent labeling technique was used to detect the presence of the major IVD ECM markers, aggrecan (ACAN), type I collagen (COL1), and type II collagen (COL2), as well as the NP-specific markers transcription factor hypoxia inducible factor 1-alpha (HIF1-α), forkhead box F1 (FOXF1), cytokeratin 19 (KRT19), and carbonic anhydrase 12 (CA12). Antibody information is summarized in **Supplementary Table 2**. Frozen sections were cut to 20 µm, washed with sodium citrate-EDTA buffer, permeabilized for 10 minutes with tris-buffered saline (TBS) with 0.3 % Triton X-100 (Fisher Scientific), and blocked with 10 % v/v goat serum in TBS for 10 minutes. Primary antibodies were applied for 1 hour at room temperature. Secondary antibodies conjugated with Alexa Fluor 647 (Molecular Probes, 1:200 dilution) were applied for 30 minutes at room. Sections were counterstained with 4’,6-diamidino-2-phenylindole dihydrochloride (DAPI) and imaged on a confocal microscope (Model A1+, Nikon Instruments Inc.). Immunofluorescent staining performed on sections without primary, secondary, or any antibodies from either mouse or rabbit species served as controls to check for non-specific staining or endogenous autofluorescence. Immunofluorescent protein expression was compared on days 0 and 14.

### 2.6 *Ex Vivo* Biomechanical Testing

#### 2.6.1 Dissection and Casting of Porcine IVDs

Lumbar spines from healthy male and female porcine donors (5 – 6 months old, 250 – 300 lb.) were purchased from Tissue Source, LLC (LaFayette, IN) and IVDs were isolated for biomechanical testing. External tissue, posterior and transverse elements were removed and individual motion segments were isolated by cutting through the midline of each vertebral body using a bone band saw (Mar-Med Inc.). Specimens were analyzed with Image J software to determine the average cross-sectional area (8.2 ± 0.4 cm^2^), potted in a polyurethane (Smooth Cast), and frozen at -20°C. IVD specimens were pre-warmed over several hours 37 °C water bath prior to testing.

#### 2.6.2 Injury, Implantation, and Biomechanical Characterization

Potted porcine IVDs (n=7 per group) were loaded on an MTS 831 Elastomer Test System using cycles of compression and tension, as has been reported prior in biomechanical studies with human [63], bovine [19, 64], and caprine IVD [65]. The motion segments were compressed to -1000 N and tensed to 100 N for 10 cycles at a rate of 0.1 Hz while maintained in a 37 °C saline bath during testing. The peak compressive loads were scaled for differences in cross-sectional area between human and porcine and selected to represent physiological pressures of jogging or climbing stairs two at a time [18]. The first nine cycles were performed as preconditioning to establish a repeatable hysteresis response and the biomechanical parameters were calculated using the 10^th^ cycle.

Each disc was subjected to the compression-tension cycles at each of the following conditions to detect changes in biomechanical parameters. First, the mechanical properties of the intact specimens were measured to obtain a baseline reference. Second, the specimens were punctured approximately 15 – 30° from the coronal plane with an 18G needle (‘‘Punctured’’ condition). Third, denucleation was performed using the needle attached to a syringe with vacuum (‘‘Denucleated’’ condition). An average of 294 ± 41 mg or 44 ± 8.9% of NP tissue was removed from the IVDs to create a cavity. Then, a compressive load from -1800 N to 0 N was applied for 50 cycles at a rate of 0.1 Hz to induce further degeneration to the disc by excessive mechanical fatigue (‘‘Degenerated’’ condition). Last, formulation S-50, cooled to 4°C, was injected into the cavity until the syringe plunger could no longer be depressed manually (‘‘Injected’’ condition). The average mass of composite hydrogel that was injected into the IVDs was 340 ± 43 mg. Implanted motion segments were incubated for 10 minutes prior to loading to allow time for complete gelation.

A MATLAB (Mathworks, Natrick, MA, USA) code was used to calculate compressive stiffness, neutral zone (NZ) stiffness, and range of motion (ROM) as described by Hom et al. 2019 [64]. Biomechanical parameters at each condition were normalized to that of the intact disc.

#### 2.6.3 Expulsion testing

Lateral bending tests were performed to observe the resistance of formulation S-50 to expulsion through the needle tract (*n* = 7). Custom-designed mechanical fixtures were created to bend the IVDs (**Supplementary Figure 1A**). Specimens were denucleated, injected with composite, and subjected to lateral bending by applying a vertical displacement (- 4 to + 4 mm) at a position 25.4 mm from the center of the specimen (**Supplementary Figure 1B)** to increase the bending angle continuously at a rate of 0.1°/s on the side opposite of the injection. The test was stopped manually when the maximum bending angle was reached due to geometric constraints of the tissue. Angles were tracked using a video camera recording at a rate of 30 frames per second. Torque was calculated as the applied force multiplied by the perpendicular distance from the axis of rotation.

#### 2.6.4 Histology of implanted porcine IVDs

Histology was performed to qualitatively assess implant conformation in the intradiscal cavity of the porcine IVD. Specimens (n=3) were either (1) intact, (2) denucleated, or (3) denucleated and injected with formulation S-50. After treatment, the discs were fixed with 4% formaldehyde in PBS for 24 h at 37 °C. Bone segments were decalcified using 5% v/v HCl in PBS for 24 h at 37 °C. Discs were embedded in frozen section compound and sectioned in the sagittal direction to 30 µm. GAGs and collagen were stained with alcian blue and picrosirius red, respectively. Cross sections were imaged using a stereoscope.

### 2.7 Statistical Analysis

Graphpad Prism 8 (San Diego, CA) was used for all statistical analyses. Welch’s t-tests were used to identify statistical differences between sample groups. All values are reported as the mean ± standard deviation (SD). Significance was set at the 95 % confidence level (p < 0.05).

## 3. Results

### 3.1. *In vitro* Material Characterization

SEM imaging was used to visualize the microscopic architecture of the hydrogel formulations (**Figure 1A**). After 14 days of swelling *in vitro*, formulation P-0 exhibited a noticeable decrease in porosity and pore diameter due to the hydrophobic behavior of the PNIPAAm macromolecular chains at 37 °C. At day 14, with the incorporation of MPs into the formulations, S-25, L-25, S-50, and L-50 qualitatively exhibited higher porosity compared to P-0, with S-50 and L-50 exhibiting the highest porosities.

**Figure 1.**
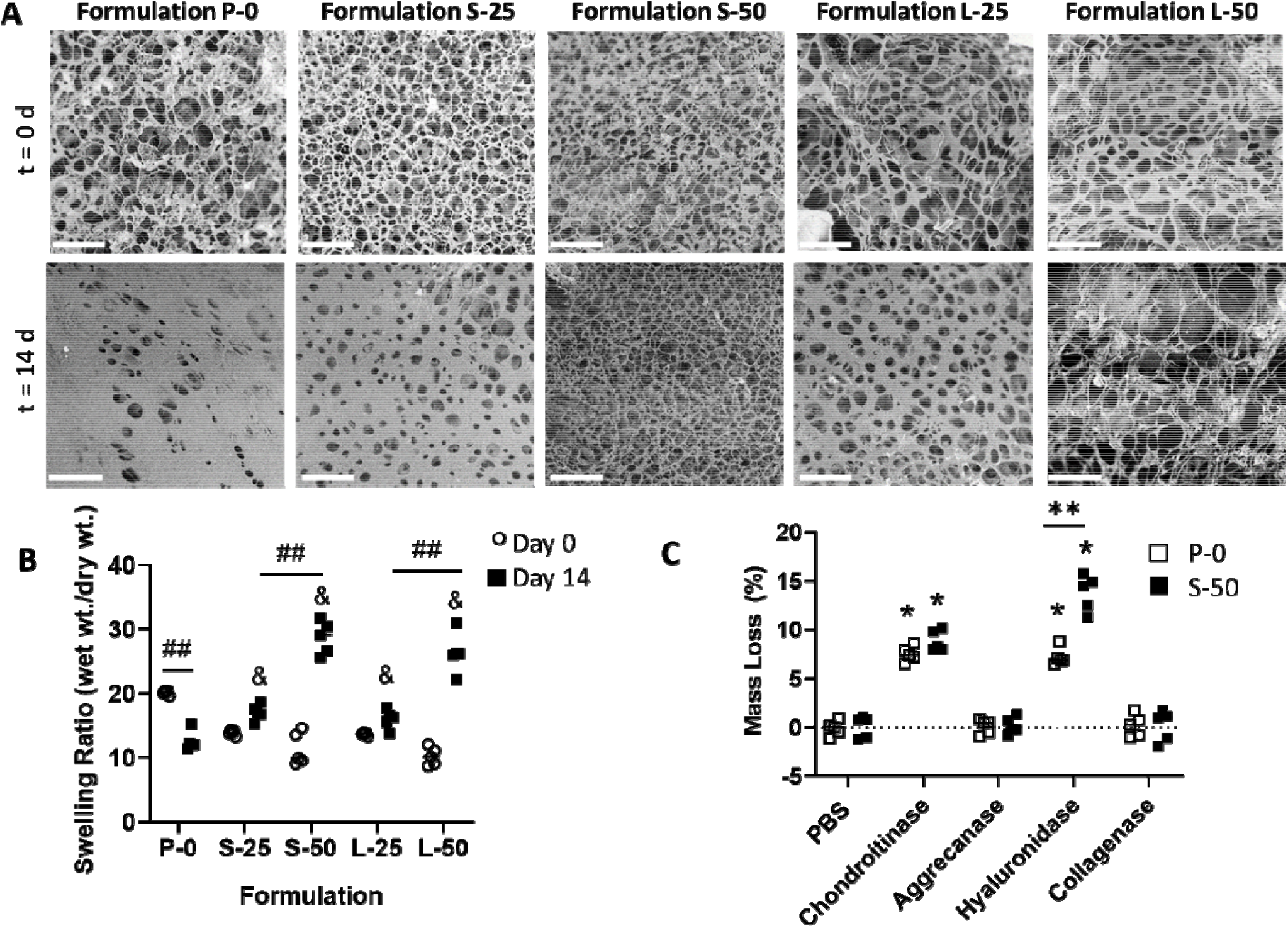
**(A)** Representative SEM images of formulations incubated in PBS at 37 °C after 0 and 14 days. Scale bars = 50 µm. **(B)** Swelling ratios of formulations incubated in PBS at 37 °C at day 0 and 14. Th ampersand (&) indicates significantly different swelling ratio compared to P-0 at day 14 (p < 0.05). The double hash symbol (##) indicates significantly different swelling ratio between two formulations or time points (p < 0.05). **(C)** Degradation behavior of formulations P-0 and S-50 at 7 days immersion in PBS or various enzymatic solutions. The asterisk (*) indicates statistically significant mass loss (p < 0.05) compared to PBS control. The double asterisks (**) indicate a significant difference in mass loss between P-0 and S-50 (p < 0.05).

The swelling ratios at days 0 and day 14 were compared for the formulations (**Figure 1B**). P-0 was the only formulation exhibiting a significant decrease (p < 0.05) in swelling ratio over this time period. The incorporation of MPs into PNIPAAM-g-CS hydrogels (S-25 and L-25) produced significant increases in swelling ratio compared to P-0 at day 14 (p < 0.01 and p < 0.05, respectively). The increase in swelling ratio compared to P-0 was more pronounced for higher concentrations of MPs (S-50 and L-50, p < 0.01). At a given concentration of MPs, varying the diameter did not significantly change the swelling ratio (p > 0.05).

The degradation behavior of formulations P-0 and S-50 in PBS and various enzymatic solutions is summarized in **Figure 1C**. No significant loss in dry mass between 0 and 7 days in PBS (p > 0.05) was measured. Exposure to the enzyme collagenase or aggrecanase did not significantly degrade the samples compared to the PBS control (p > 0.05). Compared to PBS, chondroitinase ABC caused a significant increase in mass loss of P-0 and S-50, at 7.6 ± 0.8 % and 8.9 ± 0.8 % 1.0 %, respectively (p < 0.01). Also compared to PBS, hyaluronidase caused a significant increase in mass loss for P-0 and S-50, at 7.2± 0.9% and 13.8±1.8%, respectively (p<0.01). For hyaluronidase, significantly higher mass loss was measured for S-50 compared to P-0 (p < 0.01). No other enzymes produced a significantly different mass loss for P-0 compared to S-50 (p<0.05).

#### 3.1.2 Rheological properties

A rheological temperature sweep of the formulations revealed gel points for P-0, S-25 and L-25 of 33.4 ± 0.4 °C, 30.82 ± 1.1 °C, and 32.02 ± 0.9 °C respectively (**Figure 2A**). In contrast to these formulations, gel points for S-50 and L-50 could not be identified by a crossover of G’ and G’’, due to predominantly elastic behavior over the entire temperature range (**Figure 2B**). Frequency sweeps performed at a constant temperature of 37 °C revealed viscoelastic behavior, or frequency-dependent changes for G’, G’’, and η*. Formulations P-0 (**Figure 2C,E**), S-50 (**Figure 2D,F**), as well as L-25 and L-50 (data not shown) all exhibited increases in elasticity and decreases in viscosity at higher frequency, indicated by increases in G’ and decreases in η*, respectively. Phase shift angle, δ, and G* for each of the formulations are summarized in **Table 2**. With and without MPs, values for δ were below 45° over the entire frequency range tested, another indicator that the formulations behave as viscoelastic solids under dynamic shear [66]. Compared to P-0, all formulations exhibited statistically significant increases in G* (p < 0.05), signifying a higher resistance to deformation. Regardless of diameter, increasing MP concentration from 25 to 50 mg/mL produced significant increases in G*, with S-50 exhibiting the highest G*value (p < 0.05) of all the formulations.

**Table 2.** Complex moduli (G*) and phase angle (δ) for each formulation as a function of frequency (ω). All formulations with MPs (S-25, S-50, L-25 and L-50) exhibited statistically significant increases in G* and δ compared to P-0 (p < 0.05) at both frequency levels, 0.1 and 15 Hz.

| | | $\omega$ (Hz) | |
| --- | --- | --- | --- |
| Property | Formulation | 0.1 | 15 |
| $\delta$ (°) | P-0 | $3.7 \pm 0.1$ | $1.7 \pm 0.6$ |
| | S-25 | $9.6 \pm 1.3$ | $10.7 \pm 1.5$ |
| | S-50 | $20.9 \pm 1.2$ | $21.3 \pm 1.5$ |
| | L-25 | $12.6 \pm 1.3$ | $13.6 \pm 1.7$ |
| | L-50 | $21.0 \pm 1.4$ | $26.9 \pm 1.8$ |
| $G^*$ (Pa) | P-0 | $129 \pm 20$ | $194 \pm 33$ |
| | S-25 | $268 \pm 61$ | $565 \pm 152$ |
| | S-50 | $1454 \pm 263$ | $3637 \pm 806$ |
| | L-25 | $183 \pm 39$ | $426 \pm 38$ |
| | L-50 | $814 \pm 97$ | $2215 \pm 331$ |

**Figure 2.**
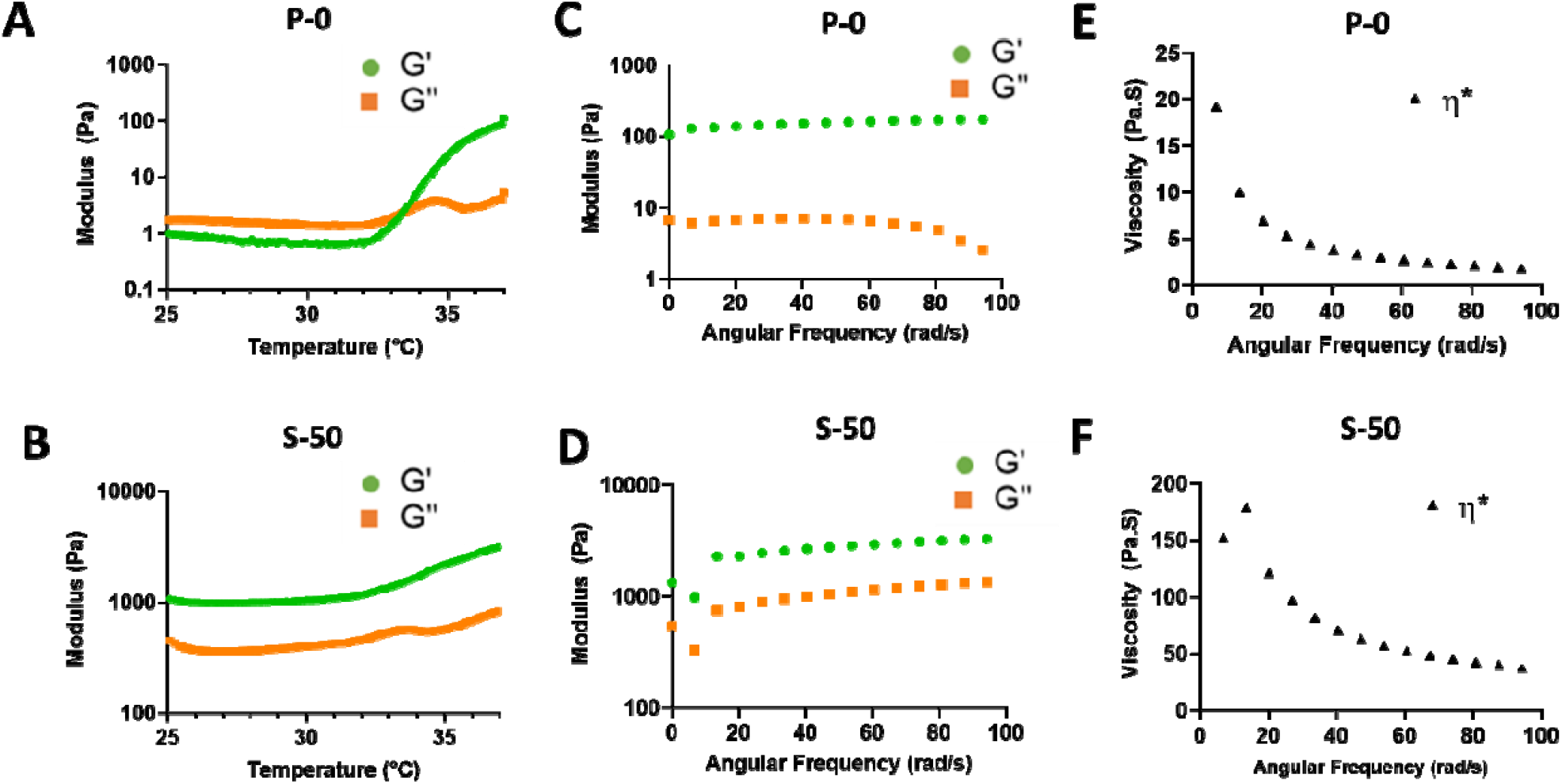
Representative rheological plots of formulations P-0 and S-50. (**A, B**) Temperature sweep from 25 to 37°C at 1 °C/min and a constant 1 % strain and 1 Hz frequency. Whereas P-0 exhibited a gel point at 33°C, identified by the G’ and G’’ crossover, S-50 did not, indicating predominantly elastic behavior over the entire temperature range due to alginate MP incorporation. (**C, D**) Frequency sweep from 0.01 to 15 Hz at a constant 1 % strain and temperature of 37 °C. Formulation S-50 exhibited higher values for G’ than P-0, signifying a higher degree of elastic behavior. **(E,F)** Frequency sweeps at 37 °C revealed a higher overall viscosity η* for S-50 than P-0, although both formulations exhibited decreasing η* at higher frequencies.

#### 3.1.3 Adhesive Properties

Histological images of the hydrogels applied to the porcine inner AF tissue substrates before adhesion testing is shown in **Figure 3**. Qualitative observation reveals that P-0 and fibrin hydrogel spread into a thin layer along the tissue surface, whereas S-50 retained its 3D shape comprised of a network alginate MPs.

**Figure 3.**
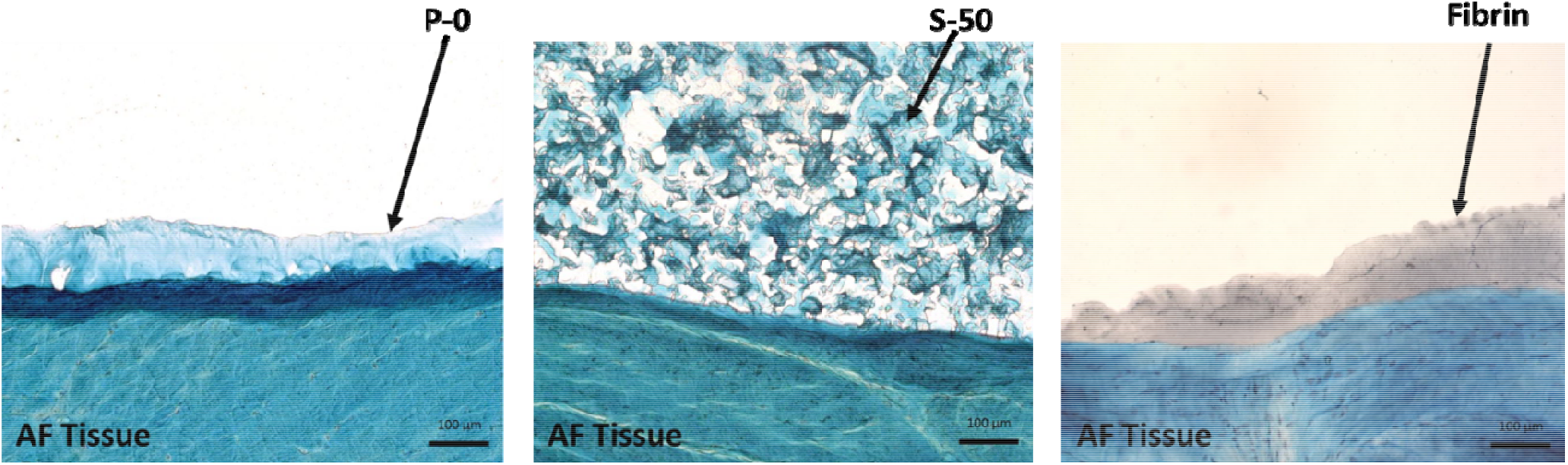
Histological staining of formulations P-0, S-50, and fibrin control applied along the porcine inner AF tissue substrate before adhesion testing. Tissue and biomaterials were stained with alcian blu and cell nuclei counterstained with Weigert’s hematoxylin. Formulation S-50, with the three-dimensional network of alginate microparticles, retained its shape when applied over the tissue surface, as opposed to P-0 and fibrin, which spread easily. Scale bars = 100 µm.

Adhesive strength to inner AF tissue was quantitated for each formulation in tension and shear. The tensile strength of fibrin was not significantly different than P-0 (**Figure 4A**, 1.83 ± 0.52 kPa for Fibrin versus 1.30 ± 0.12 kPa for P-0, p > 0.05). All of the formulations with MPs (S-25, S-50, L-25, L-50) exhibited significant increases in tensile strength compared to P-0 (p > 0.05), but only S-50 outperformed the fibrin (p < 0.01). Increasing the concentration of small MPs from 25 to 50 mg/mL produced significant increases in tensile adhesive strength (p < 0.01). However, for the large MPs, increasing concentration produced no significant changes (p > 0.05). Formulations S-50 and L-50 exhibited the highest tensile strengths of the formulations (2.79 ± 0.23 and 2.62 ± 0.53 kPa, respectively) and were not significantly different from each other (p > 0.05).

**Figure 4.**
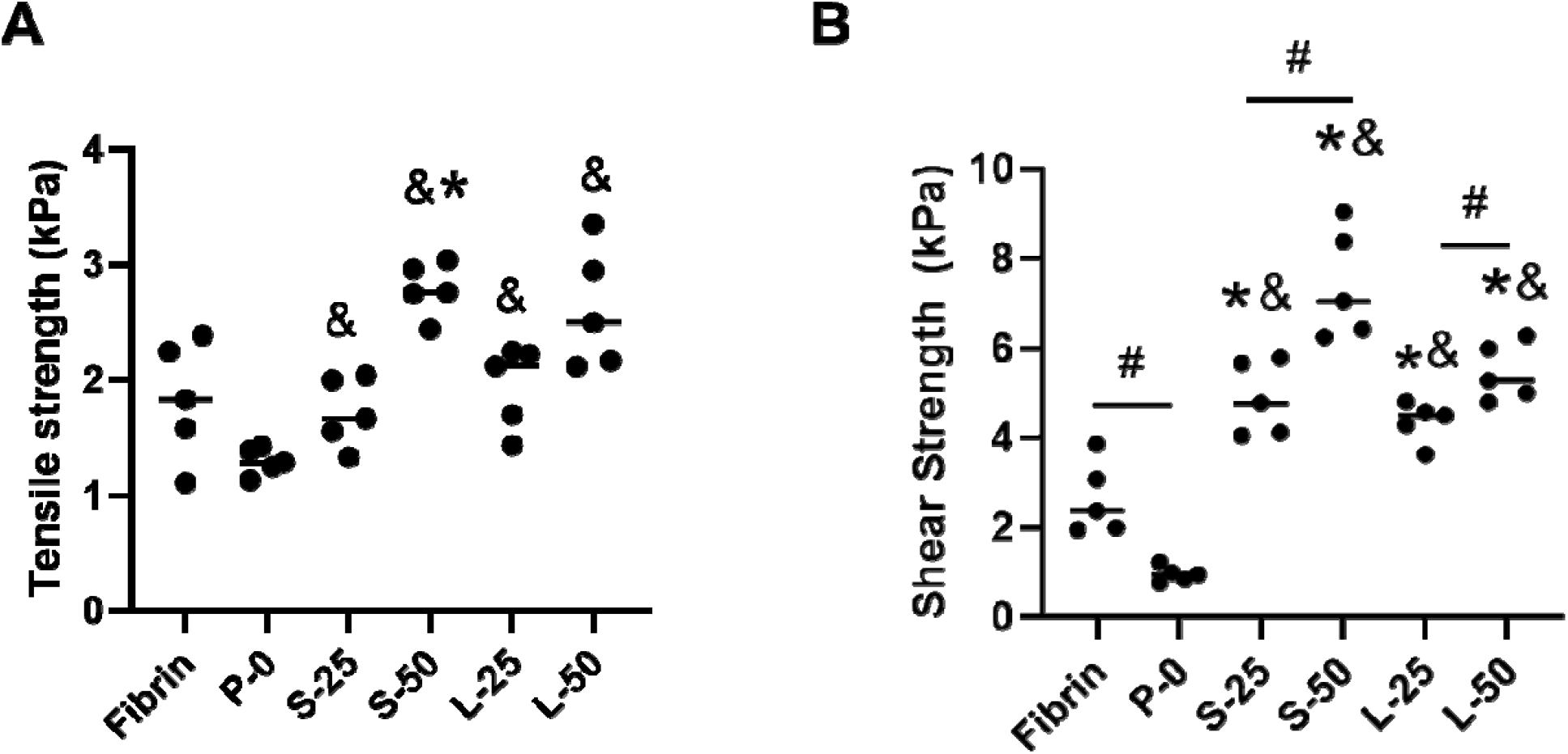
Adhesive strength of the formulations in tension **(A)** and shear **(B)** to inner AF tissue at 37 °C. An asterisk (*) indicates a statistically significant difference (p < 0.05) compared to fibrin. An ampersand (&) indicates significant difference (p < 0.05) relative to P-0. A hash symbol (#) indicates a significant difference between formulations (p < 0.05). Comparatively among all the formulations, S-50 exhibited high adhesion strength in both loading modes.

The shear strength of P-0 was significantly lower than Fibrin (**Figure 4B**, 0.96 ± 0.17 kPa versus 2.66 ± 0.81 kPa, respectively, p < 0.01). However, the shear strengths of all the formulations containing MPs (S-25, S-50, L-25, and L-50) were significantly higher than both Fibrin and P-0 (p < 0.05). Varying MP diameter did not produce any significant changes in adhesive or tensile strength (p > 0.05). However, for both MP diameters, increasing the MP concentration produced significant increases in shear strength (p < 0.02). Formulation S-50 exhibited a significantly higher shear strength compared to the other formulations (7.43 ± 1.23 kPa, p < 0.01). Overall, the formulations with MPs exhibited higher adhesive strength in shear compared to tension.

#### 3.1.4 Compressive Mechanical Properties

The compressive modulus was calculated for each formulation in unconfined and confined testing conditions (**Figure 5A and B**, respectively). Under unconfined compression, all the formulations with MPs (S-25, S-50, L-25 and L-50) outperformed P-0, with a modulus value of 1.02 ± 0.15 kPa (p < 0.01). Varying MP diameter did not produce any significant changes in unconfined compressive modulus (p > 0.05). However, for both MP diameters, increasing the MP concentration produced significant increases in unconfined compressive modulus (p < 0.01). Formulation S-50 exhibited a significantly higher unconfined compressive modulus compared to the other formulations (2.62 ± 0.14 kPa, p < 0.01).

**Figure 5.**
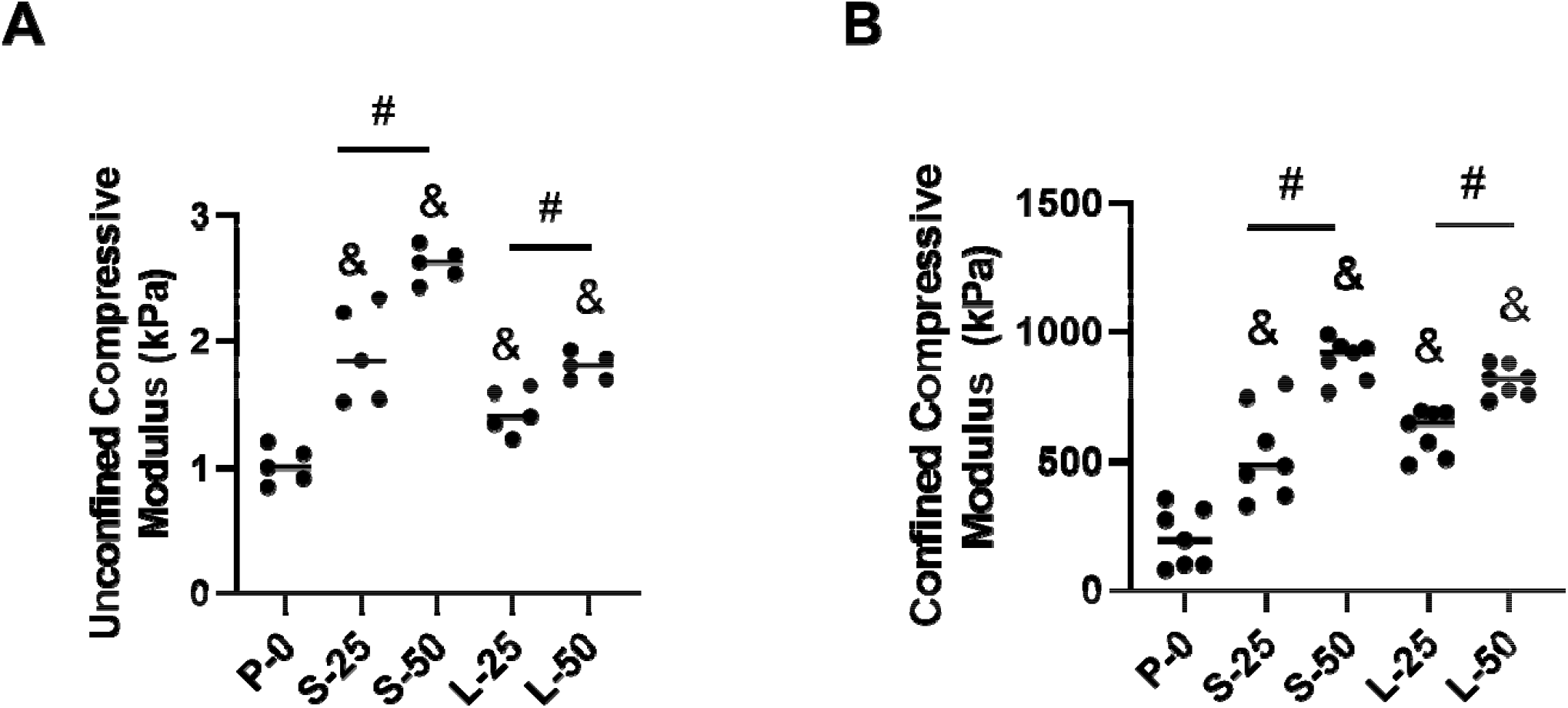
Stiffness of the formulations at 25% strain under unconfined **(A)** and confined **(B)** compression at 37 °C. An ampersand (&) indicates significant difference (p < 0.05) relative to P-0. A hash symbol (#) indicates a significant difference between formulations (p < 0.05). Comparatively among all th formulations, S-50 exhibited high compressive strength.

The confined compressive moduli for all the formulations with MPs were significantly higher than that of P-0 (p < 0.001). Varying MP diameter did not produce any significant changes in unconfined compressive modulus (p > 0.05). However, for both MP diameters, increasing the MP concentration produced significant increases in unconfined compressive modulus (p < 0.01). Formulations S-50 and L-50 had the highest confined compressive moduli (894 ± 78 kPa and 810 ± 58 kPa), but there was no significant difference between them (p = 0.05).

### 3.2 *In Vitro* Cell Culture Study

After 14 days of encapsulation, ADMSCs showed excellent cellular viability within P-0 and S-50. The proportion of living cells in P-0 (**Figure 6A**) and S-50 (**Figure 6B**) was calculated to be 91.8 ± 1.7 % and 93.4 ± 1.8 %, respectively. Both P-0 and S-50 showed significant increases in reagent reduction at day 14 relative to day 0 (p < 0.0001), indicating cell proliferation (**Figure 6C**). Reagent reduction was significantly higher for P-0 compared to S-50 at days 7 and 14 (p = 0.004, p < 0.001, respectively).

**Figure 6.**
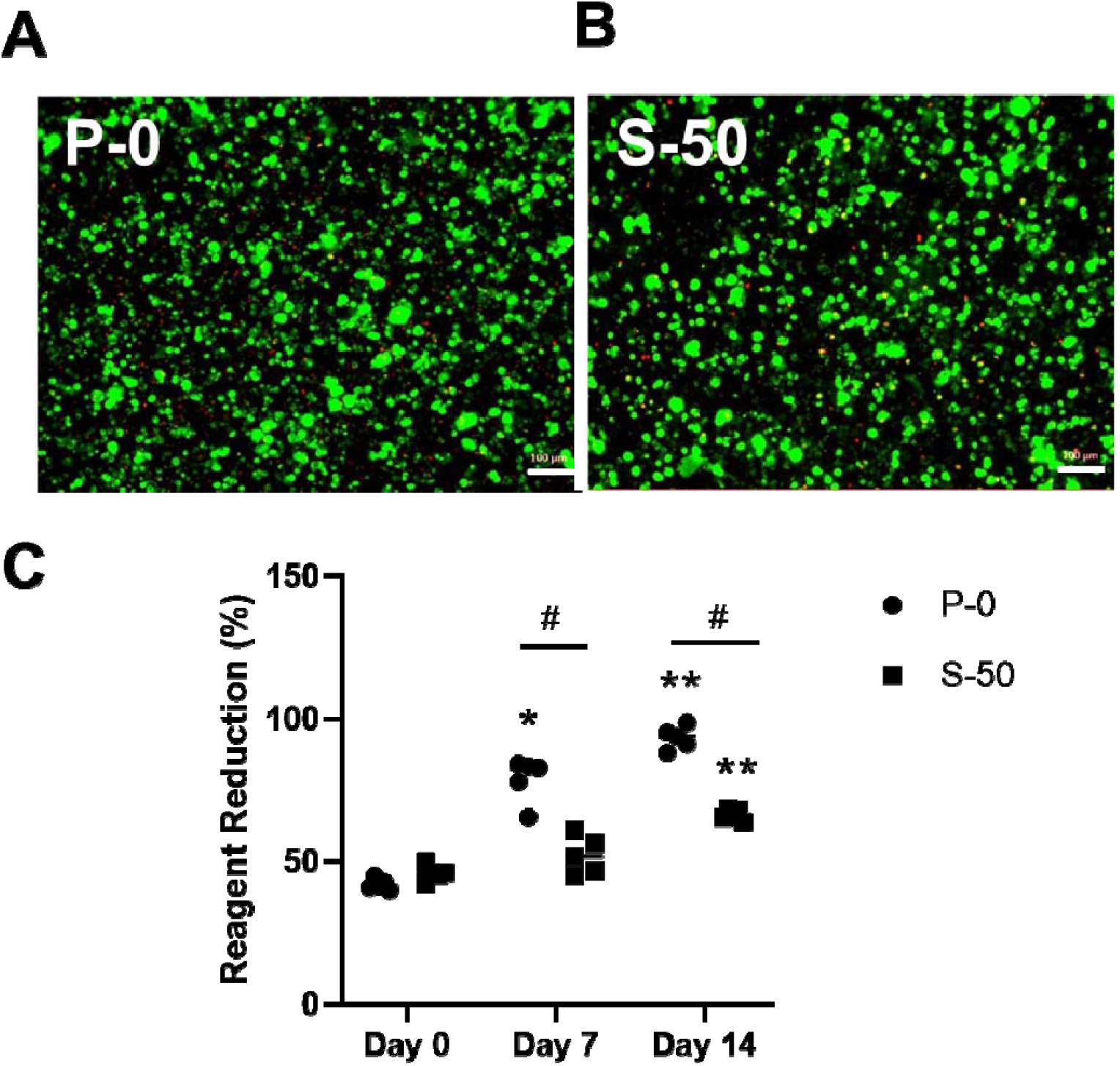
Representative Live/Dead images illustrating the viability of ADMSCs cultured for 14 day within formulation **(A)** P-0 or **(B)** S-50. Living and dead cells are shown in green and red, respectively. Scale bars = 100 µm. **(C)** Reagent reduction values calculated from alamarBlue assay results indicating th metabolic activity of ADMSCs on days 0, 7, and 14 (n=5). An asterisk (*) indicates a statisticall significant difference (p < 0.01) relative to day 0. The double asterisks (**) indicate a significant difference (p < 0.0001) relative to day 0. The hash symbol (#) indicates a significant difference (p < 0.01) between formulations P-0 and S-50.

Histological staining indicated that ADMSCs seeded in P-0 and S-50 synthesized GAGs and collagen, the major ECM molecules of NP tissue (**Figure 7**). Intensity of intracellular and extracellular staining increased for both formulations after 14 days of culture. Formulation P-0 showed relatively low deposition of GAG and collagen compared to P-0. Also, the ECM in S-50 appeared to form concentrated striations bridging gaps between encapsulated cells (**Figure 7 C,F**).

**Figure 7.**
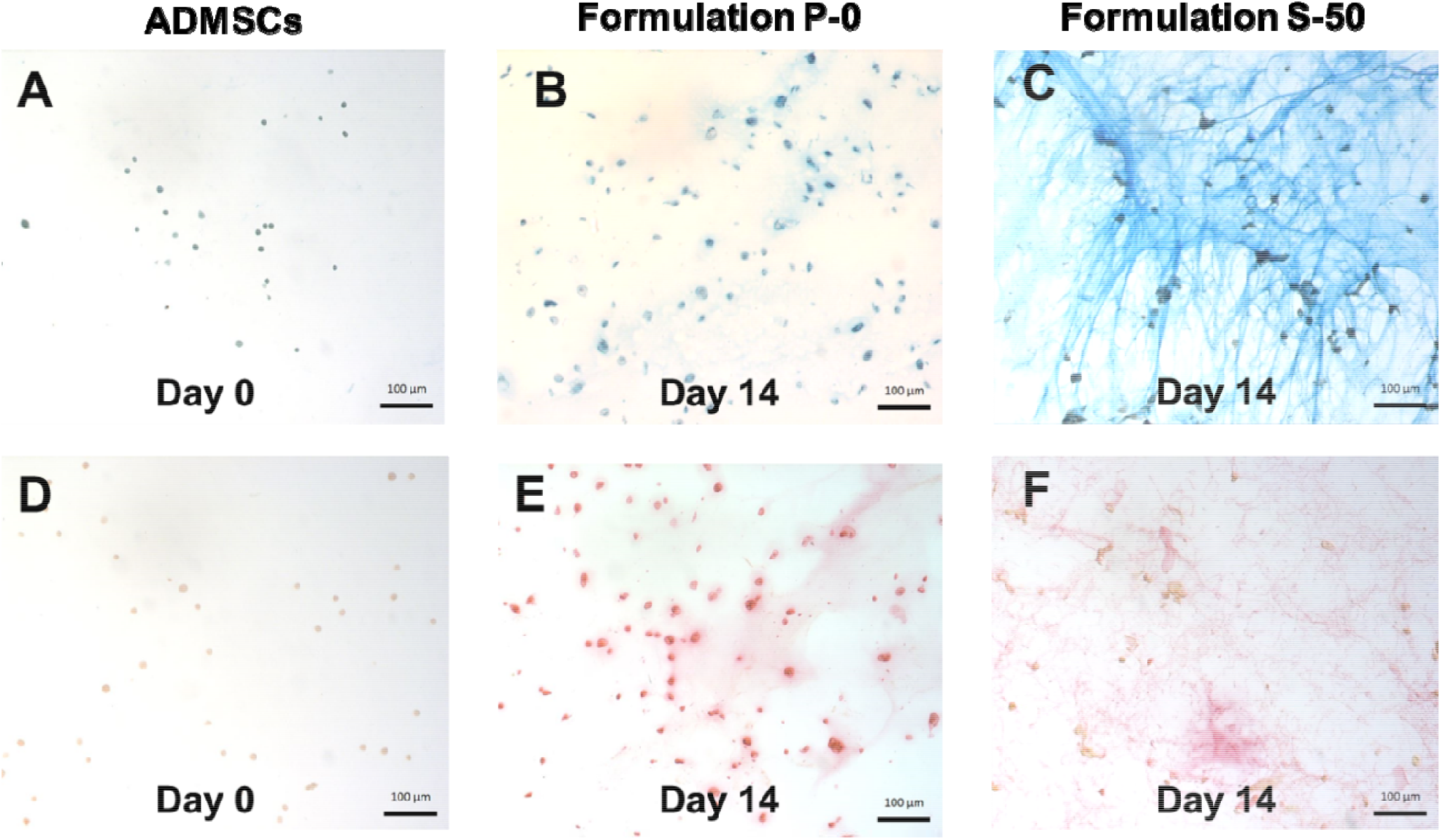
Representative histological images of ADMSCs cultured within formulation P-0 or S-50 for 0 or 14 days in the presence of soluble GDF-6. GAGs **(A-C)** and collagen **(D-F)** were stained with alcian blue and picrosirius red, respectively. Nuclei were counterstained with Weigert’s hematoxylin. Scale bars = 100 µm.

ADMSC differentiation toward an NP-like phenotype was further examined in S-50 with immunofluorescent staining. Extracellular staining of the major IVD ECM components, collagen type I, collagen type II, and aggrecan was detected after 14 days of culture (**Figure 8A-C**). Prior to culturing, undifferentiated ADMSCs showed low levels of expression for these proteins (**Figure 8D-F**). In addition, higher levels of intracellular staining for the NP-specific proteins, CA12, FOXF1, HIF1α, and KRT19 were observed at day 14 (**Supplementary Figure 1A-E**) compared to day 0 (**Supplementary Figure 1F-J**).

**Figure 8.**
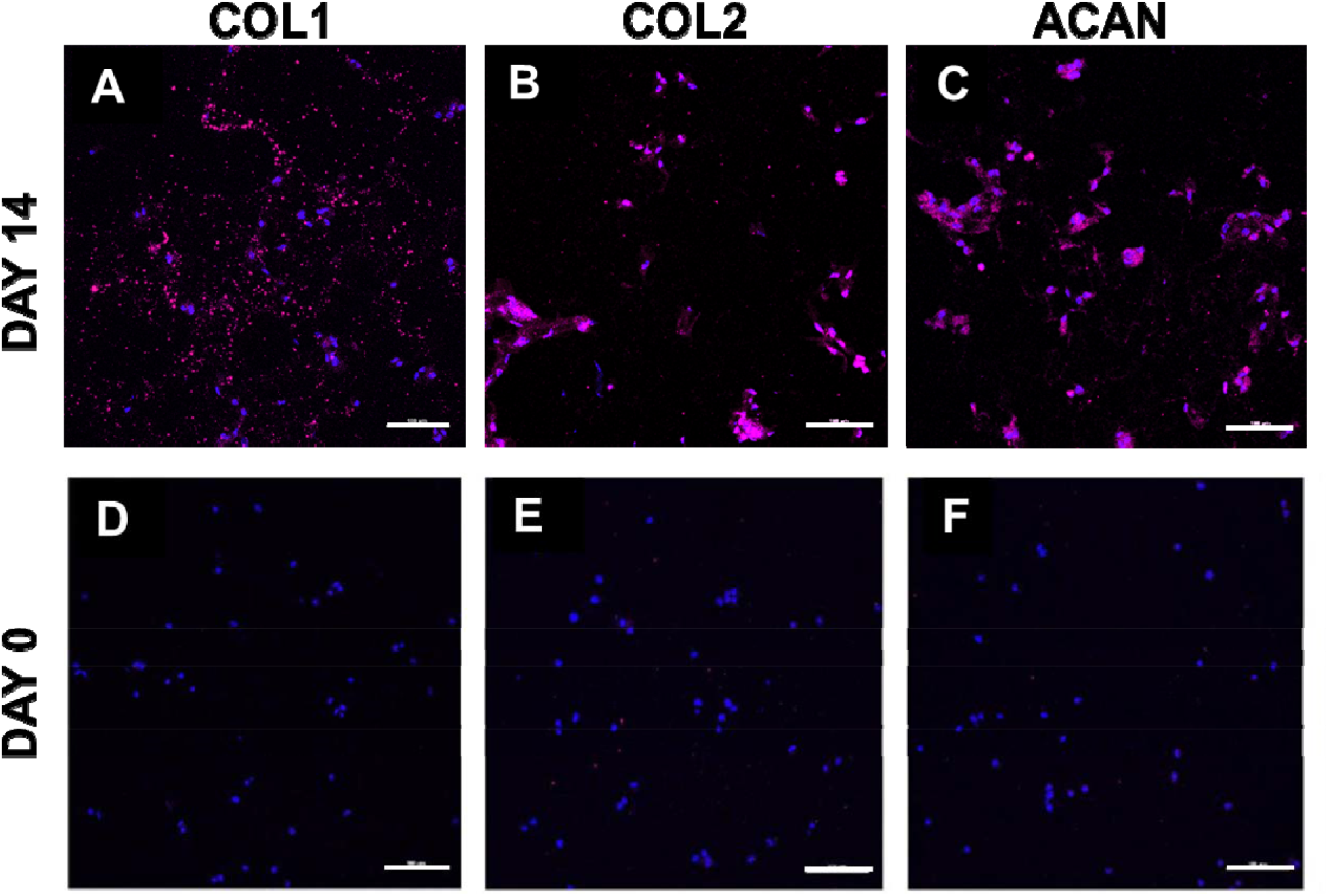
Representative immunofluorescent staining (magenta) of **(A)** COL1, **(B)** COL2, and **(C)** ACAN produced by ADMSCs cultured within formulation S-50 for 14 days in the presence of soluble GDF-6. Staining for day 0, immediately after encapsulation, is presented as a comparison in **(D-F)**. Cell nuclei are counterstained with DAPI (blue). Scale bars = 100 µm.

PCR analysis for cells encapsulated in S-50 (**Figure 9**) indicate the significant upregulation of all tested major IVD ECM and NP-specific markers (p < 0.01 for all markers relative to day 0). Among the markers, ACAN showed the highest upregulation (≈ 250-fold change, **Figure 9A**) followed by type II collagen (≈ 50-fold change, **Figure 9B**). Both type I collagen and SOX9 exhibited a relatively smaller upregulation (≈ 5-fold change, **Figure 9B and 9C, respectively**). KRT19, FOXF1, and PAX1 (**Figure 9D-F**) were the highest upregulated NP-specific markers compared to HIF1α and CA12 (**Figure 9G-H**).

**Figure 9.**
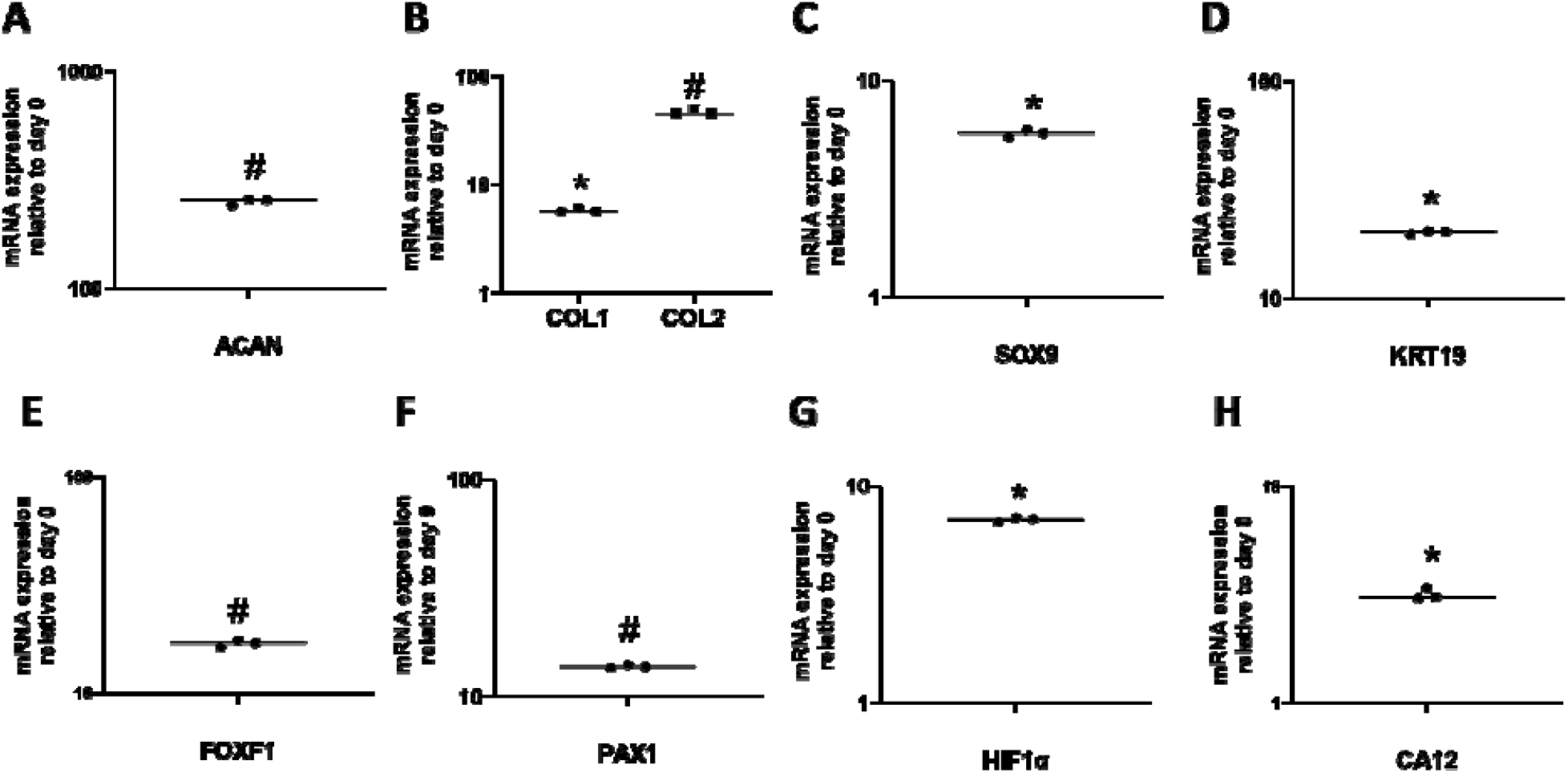
Relative gene expression profiles of ADMSCs cultured within formulation S-50 for 14 days in the presence of GDF-6. **(A)** ACAN, **(B)** COL1 and COL2, **(C)** SOX9, **(D)** KRT19, **(E)** FOXF1, **(F)** PAX1, **(G)** HIF1α, and **(H)** CA12 were upregulated relative to day 0. Data were normalized to th expression levels of GAPDH. An asterisk (*) indicates a significant upregulation (p < 0.0001) relative to day 0. The hash symbol (#) indicates a significant upregulation (p < 0.01) relative to day 0.

### *3.3 Ex Vivo* Biomechanical Testing

Formulation S-50 was also selected further evaluation in the *ex vivo* testing. The ability of the bioadhesive hydrogel to conform to surrounding disc tissue and fill an irregularly shaped defect completely was confirmed by gross observation (**Figure 10A, B**) and histology (**Figure 10C-E**).

**Figure 10.**
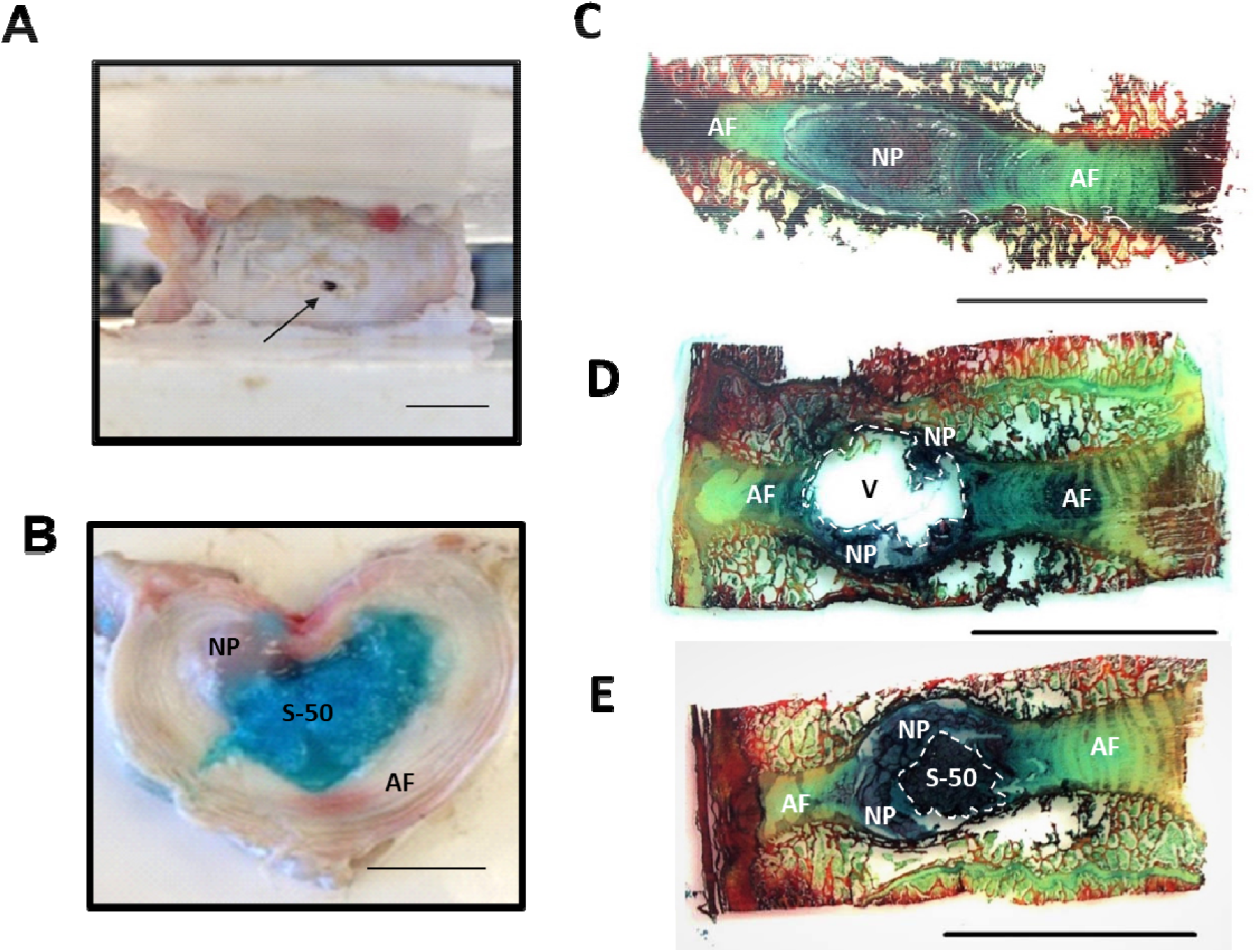
**(A)** The porcine IVD after puncture with an 18G needle. **(B)** Gross visualization of a transverse cross section of the IVD containing formulation S-50 (dyed blue) within the nuclear cavity. **(C)** Sagittal cross section of an intact IVD stained with alcian blue and picrosirius red. **(D)** A denucleated IVD. **(E)** An IVD implanted with S-50. The implant fills void space and closely interfaces with both th native NP and AF. Scale bars = 1 cm.

The axial biomechanical results are shown in **Figure 11**. Relative to intact, denucleation produced a significant decrease in NZ stiffness (p=0.01, **Figure 11A**) and an increase in compressive stiffness, though not significant (p=0.16, **Figure 11B**). The degeneration step, comprised of excessive mechanical fatigue, resulted in a statistically significant increase in compressive stiffness relative to intact (p=0.03, **Figure 11B**). The NZ and compressive stiffnesses of the injected specimens were not significantly different than that of the intact state (p=0.259 and p=0.208, **Figure 11A** and **B**, respectively). The ROM was not significantly altered from intact by injury (puncture, denucleation, degeneration) or hydrogel injection, but trended upwards with injury and downwards with implantation (**Figure 11C**). The hydrogel remained within the disc space and expulsion through the annular defect was not observed with compressive-tensile loading.

**Figure 11.**
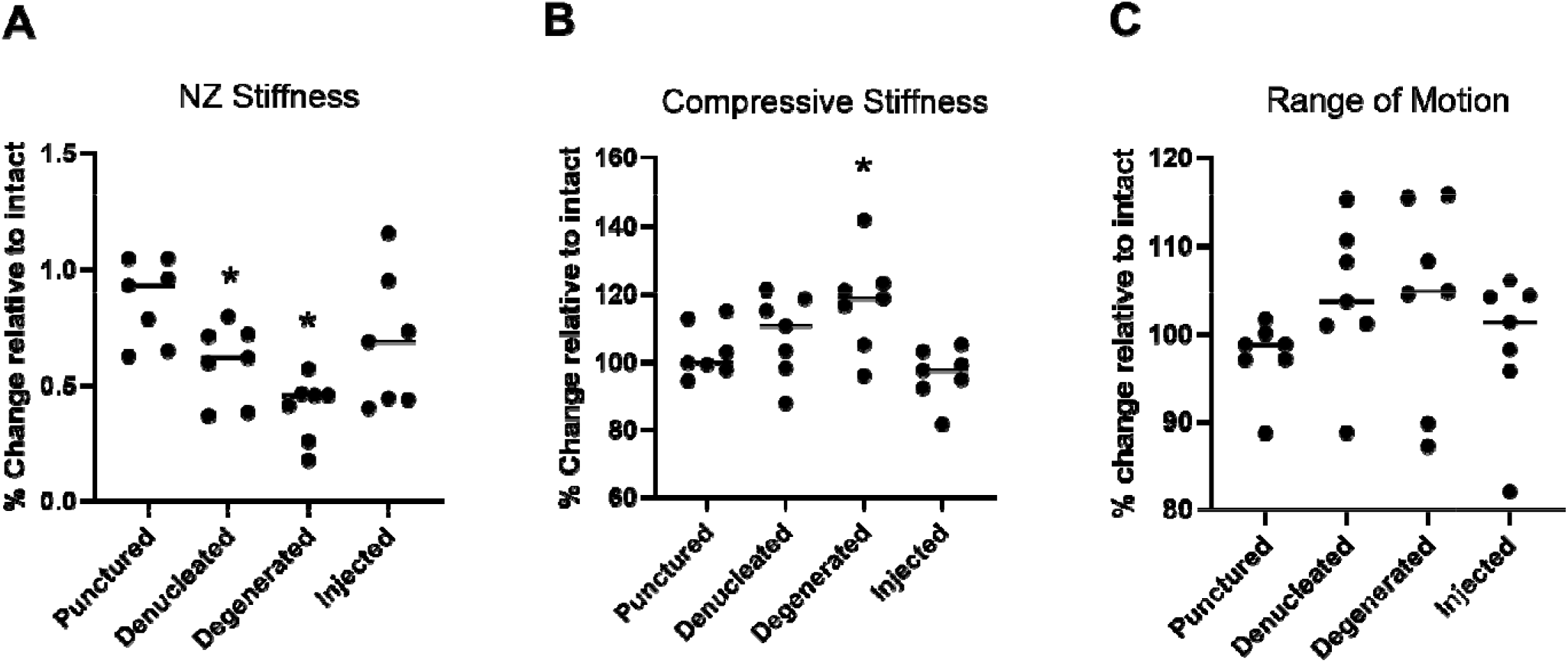
Axial biomechanical results showing **(A)** Neutral zone (NZ) stiffness, **(B)** Compressive stiffness, and **(C)** Range of motion (ROM) of bovine IVDs relative to the intact state. An asterisk (*) indicates a statistically significant change relative to intact (p < 0.05).

Lateral bending tests were performed to evaluate the composite ability to resist expulsion from within the disc space through the needle tract. Specimens were bent to an average maximum angle of 11.2 ± 1.2° (**Supplementary Figure 2C**), exhibited an average maximum torque of 5.3 ± 1.4 Nm, and showed no evidence of expulsion during testing.

## 4. Discussion

There is an important need for the development of injectable biomaterials that meet the requirements for NP replacement and repair. PNIPAAm is a promising biomaterial due to its gelation behavior between room and body temperature, but the homopolymer exhibits a low water content and poor elastic properties [67]. In previous work, we demonstrated that the polymerization of NIPAAm monomer in the presence of methacrylated CS yielded a graft copolymer (PNIPAAm-g-CS), which retained the thermosensitivity of PNIPAAm with improved water retention and compressive modulus [45]. Despite the improvements, the copolymer still exhibited water and volume loss over time, limited bioadhesive properties [46] and low solution viscosity below the LCST, characteristics identified as major obstacles to successful intradiscal implantation and biomechanical performance. Thus, in the current study, we sought to improve these properties by combining PNIPAAm-g-CS with calcium crosslinked alginate MPs to form a hydrogel composite. Structure-property relationships were investigated by varying MP size and concentration. By elucidating these relationships, we sought to also shed light on the mechanism by which MPs influence the rheological, swelling, and mechanical properties of *in situ* forming PNIPAAm-g-CS hydrogels.

In order to prepare the bioadhesive composite for this study, dry alginate MPs were suspended in aqueous solutions of PNIPAAM-g-CS immediately prior to gelation. We postulate that when the dry MPs are suspended in solution, they begin to expand as they imbibe water and packed together to form a three-dimensional ‘‘jigsaw puzzle’’ within the PNIPAAM-g-CS network. This structure, discernable in the histological image in Figure 3, imparts resistance to deformation by providing a drag force within the polymer network, an effect that is evident in multiple experimental outcomes. For instance, in the rheological study, significant increases in G* were observed for all the formulations containing MPs. Notably, the drag force increases with particle surface area. High concentrations of small MPs (S-50) induced greater increases in η* and G* compared to the same concentration of large MPs (L-50). Similarly, high concentrations of small MPs (S-50) resulted in a significant improvement in confined and unconfined compressive moduli after gelation compared to PNIPAAM-g-CS (P-0).

Alginate MPs impart tissue bonding capability to the hydrogel network. Mucoadhesion mechanisms of swellable polysaccharides have been widely reported [59, 68-70] and the principles are applicable in this system. As the alginate on the surface of the composite swells, the alginate chains become increasingly mobile and able to interact with the tissue components via hydrogen bonding, Van Der Waal forces, chain entanglement, and/or electrostatic interactions. Under tension, high concentrations of alginate MPs, whether small or large in diameter, performed equivalently, indicating that adhesive strength was primarily dependent on the amount of alginate present at the tissue interface. Likely, the drag force between particles is not induced with tensile loading. For all formulations except P-0, the magnitude of the adhesive strength was higher in shear than in tension. In shear, the flow of MP-containing hydrogel solutions into the tissue surface texture provides mechanical interlocking and a greater number of sites for bonding interactions with the tissue. This, combined with the drag force between particles, significantly improved mechanical performance of the adhesive. If the hydrogel were to expel through the AF needle tract, shear more closely mimics the mode of failure than tension, corresponding with our observation during the biomechanical studies that formulation P-0 extravasated from the porcine nuclear cavity after injection, whereas S-50 did not.

Since it outperformed the other formulations in terms of material properties, formulation S-50 was the primary focus of the *in vitro* culture experiments. Our previous studies established the biocompatibility of PNIPAAm-g-CS (formulation P-0) with encapsulated human embryonic kidney (HEK) 293 cells [45]. Clinical studies have reported improvements in Oswestry Disability Index (ODI) and Visual Analogue Scale (VAS) [71, 72] with bone marrow (BM) derived MSC injection into the IVD. However, adipose tissue, because of its relative abundance compared to bone marrow, may represent a more clinically feasible source for MSCs than bone marrow [73] and thus were selected for this study. After 14 days of culture *in vitro*, the survival and proliferation of ADMSCs encapsulated in S-50 was demonstrated with Live/Dead and alamarBlue results. Compared to P-0, ADMSCs proliferated more slowly in S-50, but nonetheless at least 90 % of ADMSCs remained viable in both formulations. We conjecture that the addition of MPs to the PNIPAAM-g-CS hydrogel imposed spatial constraints within the polymer network, limiting cell proliferation [74]. Similarly, ECM expressed by the cells appeared more striated in S-50 compared to P-0, so it is plausible that the crowding effect imposed by the MPs forced the alignment of the ECM into a more fibrous morphology. Overall, this study indicates that alginate MPs can be incorporated into PNIPAAm-g-CS networks without detrimental effects on encapsulated cells, but theoretically there is an upper limit for the concentration of MPs that can be used.

Gene expression and immunofluorescent staining of ADMSCs encapsulated in S-50 revealed the presence of several NP markers, which was expected, since GDF-6 has been reported to drive NP differentiation of ADMSCs [62]. Aggrecan gene expression was approximately 5 and 50 times higher than type II and type I collagen, respectively. Higher proportions of aggrecan to collagen (27:1) in the NP tissue of healthy adult discs has been previously reported in literature [75]. KRT19, FOXF1, and PAX1 have been recently identified as novel NP markers and were among the highest upregulated cell-related genes [76-78]. CA12 and HIF1α showed limited upregulation but are closely linked to hypoxia [79, 80], a microenvironmental condition that was not applied in this system.

Formulation S-50 was evaluated for its ability to restore the axial biomechanical behavior of a porcine IVD motion segment and resist expulsion through the needle tract in the AF. The porcine IVD has been used to model that of the human in terms of stress distributions with loading [81] and herniation behavior with flexion/extension and compression [82, 83]. Porcine IVDs have a soft nucleus pulposus, with a reported toe region modulus of 1.1 kPa [84], making it possible to denucleate through a needle attached to vacuum. Thus, we were able to induce the immediate formation of an NP cavity with minimal damage to the AF. Bovine caudal IVDs are used in currently reported *ex vivo* biomechanical studies [64, 85, 86], but the NP of this species is stiffer than that of porcine. Mechanical denucleation of bovine IVDs necessitates rongeurs, inflicting more damage to the AF than what was aimed for in this NP replacement study. Minimally invasive denucleation of bovine IVDs can be achieved by enzyme injection [5, 87-89], but digestion requires a culture period of several days with removal of vertebral body bone to preserve cell viability. Thus, porcine was chosen as the model for preliminary evaluation of biomaterial biomechanical performance, although further evaluation should be performed in other species, such as bovine.

Nucleotomy caused decreases in NZ stiffness and increases in ROM, an expected outcome since the NP plays a significant role in limiting axial deformation under low loads [13, 90, 91]. Compressive stiffness of the motion segments, a parameter measured at high loads, trended upwards with nucleotomy and mechanical degeneration as a result of the transfer of load to the stiffer IVD components [19]. Hydrogel implantation restored these biomechanical parameters to the intact state, a promising preliminary indication of the functional behavior of the composite. Yet, the hydrogel design still needs optimization for clinical translation. For instance, the mechanical properties of PNIPAAm-g-CS + MPs likely need to be increased to overlap with levels of the human. Formulation S-50, with an average unconfined compressive modulus of 2.7 kPa, is weaker than native NP tissue, ranging from 3 – 5 kPa [92]. The same formulation exhibited an average confined compressive modulus of 893 kPa, only approaching the native NP tissue value of 1 MPa [93]. Last, with complex moduli *G\** between 1.4 to 3.5 kPa, S-50 fell short of mimicking the G* of the native NP in the same frequency range, 7.4 to 19.8 kPa [66].

Another important consideration is that the material behavior of PNIPAAm-g-CS + MPs is likely to change over time. Water is known to act as a plasticizer in hydrogels [94], but the swelling kinetics of PNIPAAM-g-CS + MPs *in situ* will depend on osmotic pressure of the surrounding tissues [95]. Alginate dissolution will induce a loss of mechanical reinforcement and adhesion strength, but the rate at which this occurs depends on the ion concentration in the milieu surrounding the biomaterial [96]. Simultaneously, encapsulated cells will remodel the hydrogel network and secrete ECM [97, 98], also impacting hydrogel properties over time. Human or bovine IVD organ culture models [99-101] are the most appropriate tools for ascertaining long term hydrogel behavior within the context of an IVD-mimetic osmotic pressure, biochemical composition, and biomolecular microenvironment. While such studies are out of the scope of the current work, it is exciting to note that the two phases in the PNIPAAm-g-CS + MP composite system can be modified to tune short and long-term behavior. For the MP phase, increasing alginate concentration would slow MP dissolution [59], prolonging mechanical performance and bioadhesive interactions with the tissue. Another option to improve the long term bioadhesive stability of the system is to employ a recently reported two-part repair strategy [102], where a chemically-functionalized polymer layer would be placed between the bulk phase (in this case, PNIPAAm-g-CS) and surrounding AF, covalently linking the bulk phase to the tissue interface.

Despite the need for continued development, we posit that we have developed a useful platform for IVD tissue engineering. The concept of encapsulating re-hydrating polysaccharide-based MPs within a hydrogel structure can have important utility beyond the scope of this study, such as the controlled delivery of bioactive molecules for improving regenerative outcomes. From a broader perspective, we posit that the concept can be applied for improving the properties of *in situ* forming cell carriers in a variety of regenerative orthopaedic applications.

## 5. Conclusion

The inclusion of alginate MPs within PNIPAAm-g-CS networks is an effective method of increasing initial injectability, bioadhesive interactions, and mechanical performance. Gene expression, histology and immunohistochemistry results indicate that networks comprised of PNIPAAM-g-CS + alginate MPs supports differentiation to an NP phenotype. When implanted *ex vivo* into the intradiscal cavity of degenerated porcine IVDs, PNIPAAm-g-CS + alginate MPs restores the compressive and NZ stiffnesses to intact values. The composite also resists expulsion under tension-compression and lateral bending. Based on these results, we conclude that PNIPAAm-g-CS + alginate MPs has promise as an injectable system for NP replacement and regeneration and warrants further investigation.

## Acknowledgements

The work in this publication was supported by the National Institute of Arthritis and Musculoskeletal and Skin Diseases and the National Institute of Biomedical Imaging and Bioengineering of the National Institutes of Health under Award Number 1R15 AR 063920-01 and by the New Jersey Health Foundation under Award Number PC-2316. The content is solely the responsibility of the authors and does not necessarily represent the official views of the National Institutes of Health or the New Jersey Health Foundation.

## Authors’ Contributions

T.C, K.D., J.K., C.I, and A.J.V. designed the research. T.C. performed the experiments. A.J.V. supplied the materials. T.C., A.J.V., J.K., C.I., and K.M. analyzed the data. A.J.V. and T.C. wrote the manuscript and all authors revised.

## Data Availability Statement

The processed data required to reproduce these findings are available by contacting the authors.

---

**Supplementary Figure 1.**
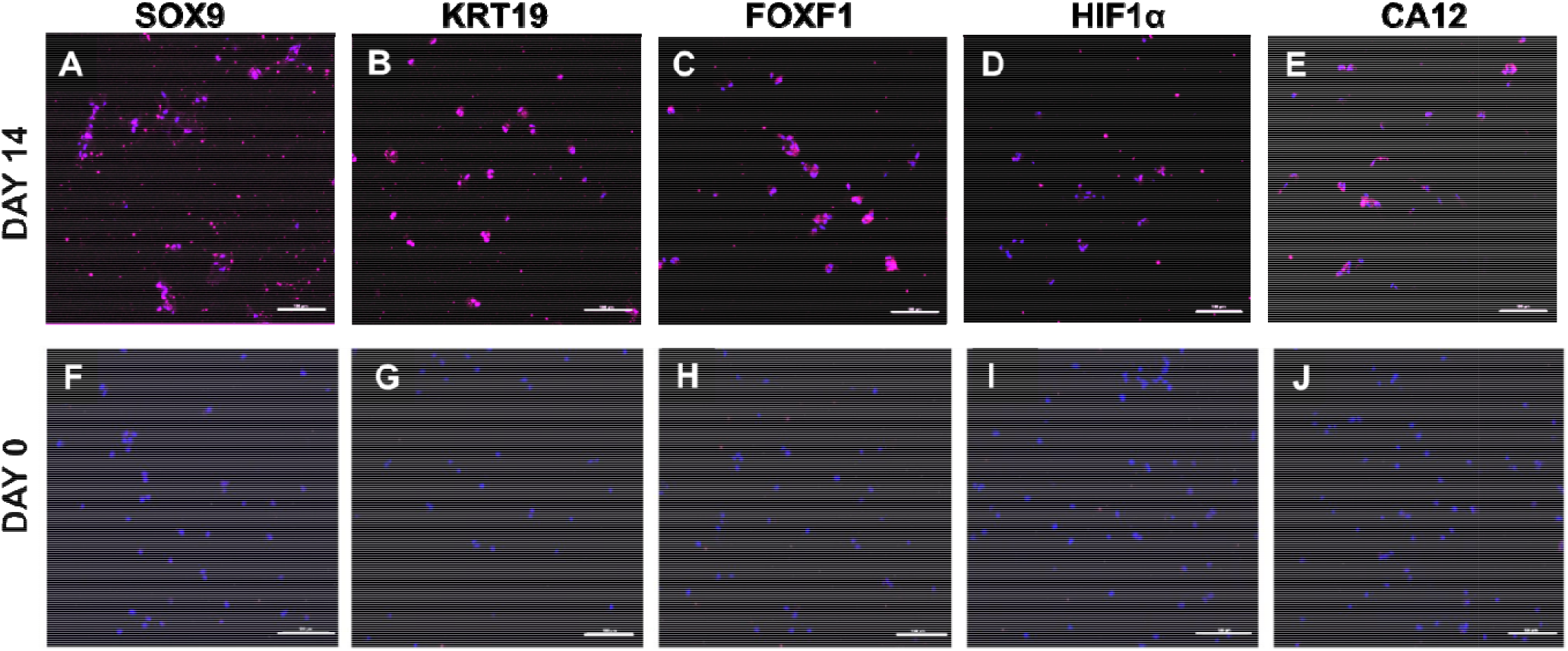
Representative immunofluorescent staining (magenta) of **(A)** SOX9, **(B)** KRT19, **(C)** FOXF1, **(D)** HIF1α, and **(E)** CA12 produced by ADMSCs cultured within S-50 for 14 day in the presence of soluble GDF-6. Staining for day 0, immediately after encapsulation within th bioadhesive, is presented as a comparison in **(F-J)**. Cell nuclei are counterstained with DAPI (blue). Scale bars = 100 µm.

**Supplementary Figure 2.**
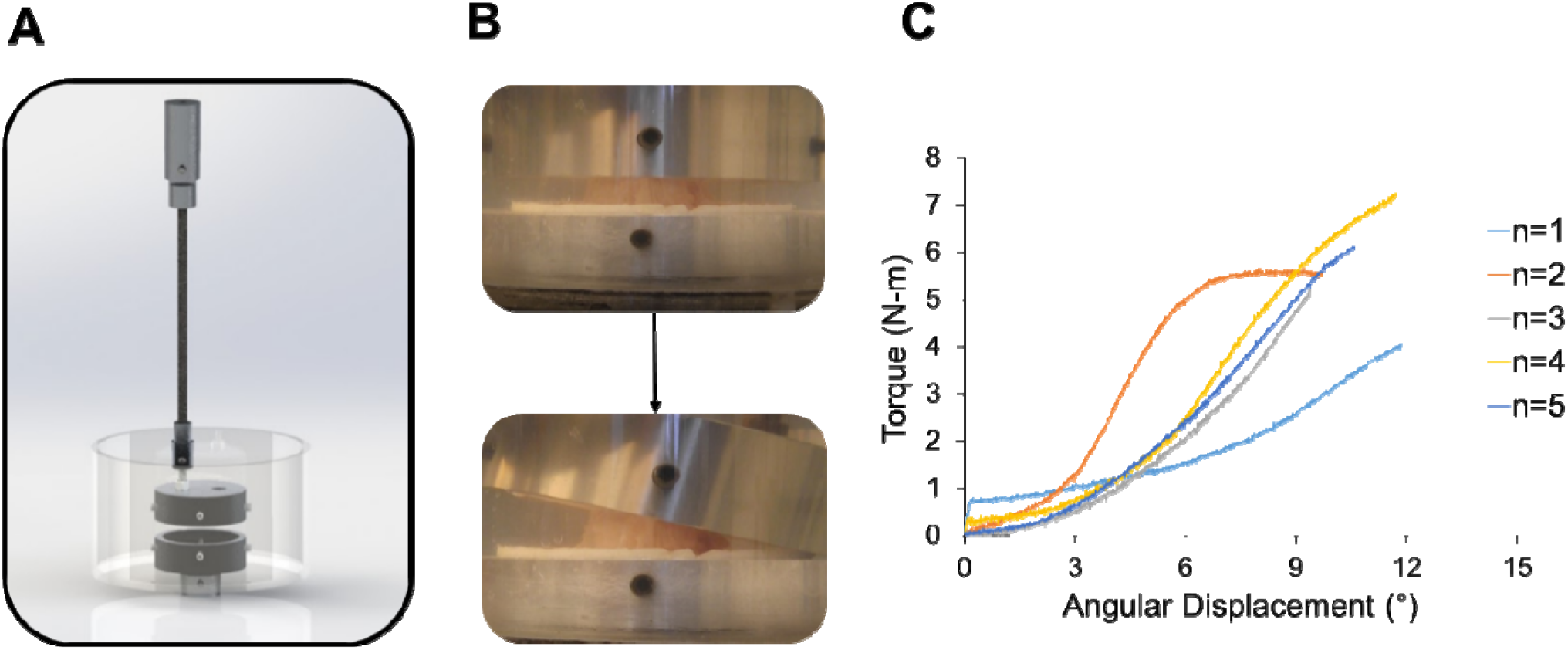
**(A)** Custom-made mechanical fixtures designed to induce lateral bending of the IVD specimen. The vertical rod is offset 25.4 mm from the center of the stainless steel cup and affixed to a freely-rotating hinge allowing for rotational movement. **(B)** High magnitude extrusion test where th angle was continuously increased at a rate of 0.1°/sec on the side opposite to the injection site. The test was stopped manually when the maximum bending angle was reached due to geometric constraints of the tissue. **(C)** Torque versus angular displacement curves for n=5 repeats of the high magnitude extrusion test. The specimens were compressed to average maximum angle of 11.2 ± 1.2° with no evidence of herniation.

**Supplementary Table 1.**
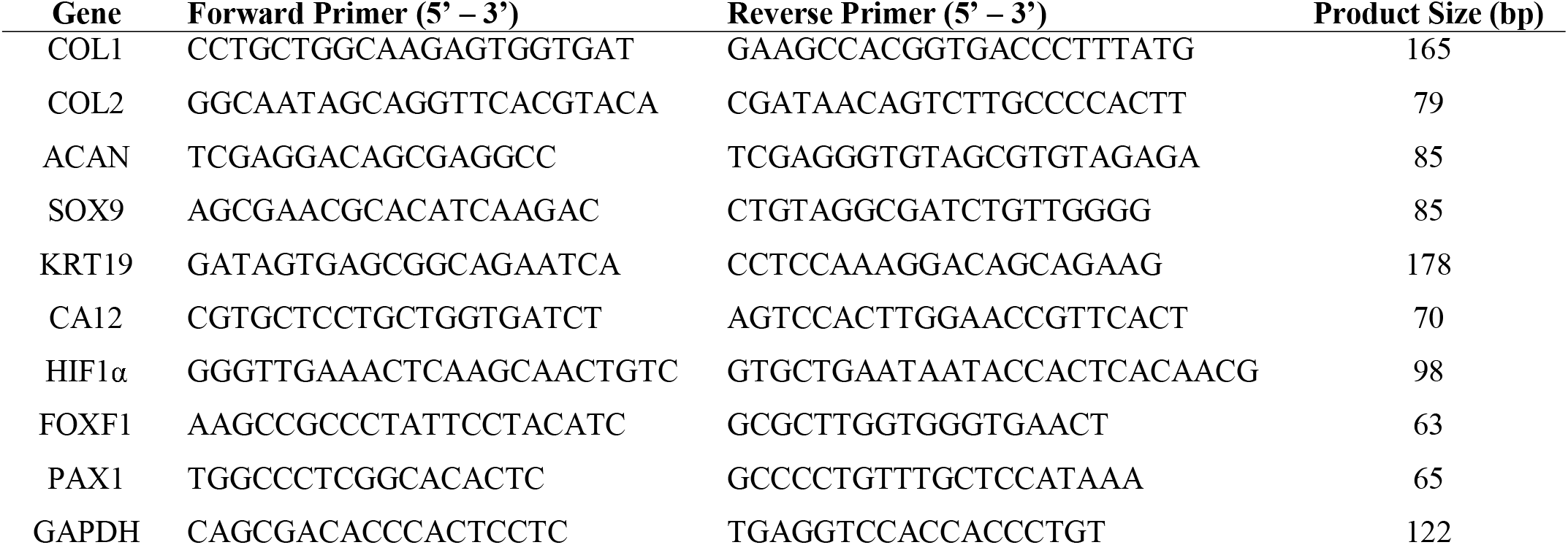
Genes of interest for ADMSCs cultured within formulation S-50 for 14 days in the presence of soluble GDF-6.

**Supplementary Table 2.** Proteins of interest for immunofluorescent labeling of ADMSCs cultured within formulation S-50 for 14 days in the presence of soluble GDF-6.

| **Protein** | **Manufacturer** | **Antibody Type** | **Species** | **Clonality** | **Dilution** |
| --- | --- | --- | --- | --- | --- |
| COL1 | ab90395 | Primary | Mouse anti-human | Monoclonal | 1:100 |
| COL2 | ab185430 | Primary | Mouse anti-human | Monoclonal | 1:200 |
| ACAN | ab3778 | Primary | Mouse anti-human | Monoclonal | 1:50 |
| SOX9 | ab76997 | Primary | Mouse anti-human | Monoclonal | 1:100 |
| KRT19 | ab7754 | Primary | Mouse anti-human | Monoclonal | 1:200 |
| CA12 | ab195233 | Primary | Rabbit anti-human | Monoclonal | 1:50 |
| HIF1α | ab51608 | Primary | Rabbit anti-human | Monoclonal | 1:100 |
| FOXF1 | ab168383 | Primary | Rabbit anti-human | Monoclonal | 1:100 |

